# A novel SHAPE reagent enables the analysis of RNA structure in living cells with unprecedented accuracy

**DOI:** 10.1101/2020.08.31.274761

**Authors:** Tycho Marinus, Adam B. Fessler, Craig A. Ogle, Danny Incarnato

**Author notes:** **To whom correspondence should be addressed:** Danny Incarnato. These authors equally contributed to this work (Tycho Marinus, Adam B. Fessler).

## Abstract

Due to the mounting evidence that RNA structure plays a critical role in regulating almost any physiological as well as pathological process, being able to accurately define the folding of RNA molecules within living cells has become a crucial need. We introduce here 2-aminopyridine-3-carboxylic acid imidazolide (2A3), as a general probe for the interrogation of RNA structures *in vivo*. 2A3 shows moderate improvements with respect to the state-of-the-art SHAPE reagent NAI on naked RNA under *in vitro* conditions, but it significantly outperforms NAI when probing RNA structure *in vivo*, particularly in bacteria, underlining its increased ability to permeate biological membranes. When used as a restraint to drive RNA structure prediction, data derived by SHAPE-MaP with 2A3 yields more accurate predictions than NAI-derived data. Due to its extreme efficiency and accuracy, we can anticipate that 2A3 will rapidly take over conventional SHAPE reagents for probing RNA structures both *in vitro* and *in vivo*.

## INTRODUCTION

RNA structure is a jack-of-all-trades. Besides being crucial to the function of structural non-coding RNAs (ncRNAs), RNA structural elements are now widely recognized as key players in almost any essential biological process, ranging from transcriptional and post-transcriptional control of gene expression, to catalysis and sensing of environmental stimuli such as metabolite and temperature changes (1).

In light of its importance, being able to define the structure of an RNA molecule is a key need towards understanding its mechanism of action. It is however still impossible for most RNAs to define their secondary structures starting solely from their primary sequence, a dilemma dubbed *RNA folding problem* by Prof. Herschlag in 1995 (2). Main reason for this is the huge size of the RNA folding space, so that a *n* nucleotides-long RNA can theoretically fold in up to 1.8^*n*^ distinct conformations (3).

A widely used approach to define the putative structure of an RNA based on its sequence relies on predicting its minimum free energy (MFE) structure, the theoretically most stable structure, based on a set of experimentally-determined thermodynamic parameters known as *nearest neighbor model* (or Turner rules) (4). Unfortunately, the accuracy of MFE predictions is very limited, with only ∼66% of predicted base pairs actually occurring in experimentally-validated structures (5). This is likely to be a consequence of the fact that, beyond thermodynamics, a multitude of cellular factors play a key role in regulating RNA folding in the context of the living cell, such as RNA post-transcriptional modifications, protein binding, crowding, and many others (6–8). The accuracy of thermodynamics-driven RNA structure determination algorithms can be however improved by incorporating experimental constraints from RNA footprinting experiments. These experiments are based on the use of specific chemicals or nucleases to probe the conformation of RNA residues. When informed with these experimentally-derived constraints, the accuracy of RNA structure prediction algorithms greatly improves (9–13).

Only few chemical probes can readily permeate cell membranes and can thus be used *in vivo*, while the majority of them are only suitable for *in vitro* and *ex vivo* RNA structure probing experiments. Two classes of compounds are mainly used to date. On one hand, nucleobase-specific probes, such as dimethyl sulfate (DMS) and 1-ethyl-3-(3-dimethylaminopropyl)carbodiimide (EDC), can be respectively used to probe adenine (A) and cytosine (C), or guanine (G) and uracil (U) bases (14, 15). On the other hand, selective 2’-hydroxyl acylation analyzed by primer extension (SHAPE) reagents can be used to probe the local flexibility of the RNA backbone in a nucleobase-independent fashion (16), as they can form adducts with the 2’-OH of the ribose moiety. Among SHAPE reagents, 3 have been extensively characterized: 1-Methyl-7-nitroisatoic anhydride (1M7), 2-methyl-3-furoic acid imidazolide (FAI) and 2-methylnicotinic acid imidazolide (NAI) (17, 18), with the latter being the most effective at probing RNA in living cells (19, 20).

Typical readout of RNA probing experiments is the detection of reverse transcription (RT) drop-off events induced by the probing reagent-modified RNA bases (21). More recently, it has been shown that, by either using specific RT enzymes or RT conditions, it is possible to favor the RT read-through on modified RNA bases, ultimately resulting into the recording of multiple modification sites as mutations within the same cDNA molecule. These techniques have been therefore dubbed mutational profiling approaches (MaP) (22–25) and are rapidly taking over traditional RT drop-off methods.

While the use of SHAPE compounds over nucleobase-specific probes has in theory a considerable advantage, as they enable the unbiased probing of all four RNA bases, reagents such as DMS still exhibit a higher signal-to-noise ratio in MaP experiments. To this end, we here sought to optimize *in vivo* RNA probing by synthesizing six new SHAPE reagents and testing them in Gram-positives, Gram-negatives and mammalian cells. One of these compounds, 2-aminopyridine-3-carboxylic acid imidazolide (2A3), showed both increased reactivity with RNA and higher permeability to biological membranes, resulting in a significantly higher signal-to-noise ratio when compared to NAI. Importantly, when used to perform experimentally-informed prediction of *in vivo* RNA structures, 2A3 produced markedly more accurate predictions than NAI.

## MATERIALS AND METHODS

### Synthesis of SHAPE reagents

For the design of novel SHAPE reagents, we first compiled a list of commercially-available carboxylic acids. We then selected those containing nitrogen groups or for which the putative adduct-forming group had a pKa close to (or slightly above) 7. The pKa values in water were obtained (where possible) using the DataWarrior tool (26). The underlying rationale was to slow-down the hydrolysis is water and to favor reaction with RNA. For the synthesis of SHAPE reagents, 2-methylpyridine-3-carboxylic acid (cat. 325228), isoquinoline-6-carboxylic acid (cat. APO455831358-250MG), indoline-5-carboxylic acid (cat. CDS019637), 1-methylimidazole-4-carboxylic acid (cat. 679720), 6-aminopyridine-3-carboxylic acid (cat. 216879), benzotriazole-5-carboxylic acid (cat. 304239), nicotinic acid (cat. N4126), 2-aminopyridine-3-carboxylic acid (cat. A68300) and 1,1′-Carbonyldiimidazole (CDI, cat. 21860) were purchased from Sigma Aldrich. For the synthesis of NAI, I6, I5, 1M4, B5, 6A3, NIC and 2A3, respectively 137.14 mg, 173.17 mg, 163.17 mg, 126.11 mg, 163.13 mg, 138.12 mg, 123.11 mg and 138.12 mg were resuspended in 500 μl DMSO anhydrous (Sigma Aldrich, cat. 276855). ∼1.3 g of CDI were then resuspended in 4 mL DMSO anhydrous and 500 μl of this solution were added to each 500 μl of carboxylic acids in DMSO while constantly stirring, over a period of 5 minutes. Reaction mixtures were then incubated at room temperature with constant stirring for 1 hour. Residual unreacted material was then removed by briefly centrifuging the solutions for 5 minutes at 17,000 x *g*, after which the cleared solution was transferred to a clean tube, aliquoted into 50 μl aliquots and stored at −80°C.

### Chemical characterization

NMR spectra were recorded on a JEOL-ECA 500 MHz NMR spectrometer or JEOL-ECX 300 MHz NMR in DMSO-d_6_ or D_2_O. Spectra were referenced to the respective solvent peaks for ^1^H and ^13^C. Routine ESI MS data were collected on a ThermoFisher MSQ Plus single quadrupole mass spectrometer. High resolution MS data were analyzed on a ThermoFisher Orbitrap XL in positive mode. Chemical characterization is detailed in the Supplementary Information.

### Half-life determination

SHAPE electrophiles were dissolved in 700 μL PBS buffer 1X pH 7.4 prepared with D_2_O, and any undissolved solids were centrifuged out prior to transfer to the NMR tube. As needed, some samples were dissolved in DMSO d-6 (10% final concentration) and added to D_2_O to achieve quantifiable concentrations. Samples were incubated at 25°C or 37°C and kinetic ^1^H-NMR scanning was performed. The disappearance of product resonances and reappearance of starting materials (correlating to hydrolysis) was monitored as previously described (27). Half-life data is provided in Supplementary Table S1.

### Extraction of native deproteinized *E. coli* rRNAs

Deproteinized *E. coli* RNA was prepared essentially as previously described (24), with minor changes. Briefly, a single colony of *Escherichia coli* K-12 DH10B was picked and inoculated in LB medium without antibiotics, then grown to exponential phase (OD_600_ ∼ 0.5). 2 mL aliquots were collected by centrifugation at 1000 x *g* (4°C) for 5 min. Cell pellets were resuspended in 1 mL of Resuspension Buffer [15 mM Tris-HCl pH 8.0; 450 mM Sucrose; 8 mM EDTA], and lysozyme was added to a final concentration of 100 μg/mL. After incubation at 22°C for 5 min and on ice for 10 min, protoplasts were collected by centrifugation at 5000 x *g* (4°C) for 5 min. Pellets were resuspended in 120 μl Protoplast Lysis Buffer [50 mM HEPES pH 8.0; 200 mM NaCl; 5 mM MgCl_2_; 1.5% SDS], supplemented with 0.2 μg/μl Proteinase K. Samples were incubated for 5 min at 22°C and for 5 min on ice. Sample was then extracted 2 times with phenol:chloroform:isoamyl alcohol (25:24:1, pre-equilibrated 3 times with RNA folding buffer [50 mM HEPES pH 8.0; 200 mM NaCl; 5 mM MgCl_2_]), and once with chloroform. 20 U SUPERase•In™ RNase Inhibitor (ThermoFisher Scientific, cat. AM2696) were then added and RNA was equilibrated at 37°C for 20 min prior to probing.

### *Ex vivo* probing of *E. coli* rRNAs

180 μl of deproteinized *E. coli* rRNAs in RNA folding buffer, pre-equilibrated at 37°C for 20 min, were mixed with 20 μl of each SHAPE compound (assuming complete conversion, stocks should be ∼1 M, resulting in a final concentration of 100 mM in the probing reaction). RNA was then allowed to react at 37°C for 15 minutes, with moderate shaking, after which 200 μl of 1 M DTT were added, to quench the reaction. Samples were then vortexed briefly and 1 mL of ice-cold QIAzol Lysis Reagent (Qiagen, cat. 79306) was added to each sample, followed by extensive vortexing.

### *In vivo* probing of *E. coli* cells

A single colony of *Escherichia coli* K-12 DH10B was picked and inoculated in LB medium without antibiotics, then grown to exponential phase (OD_600_ ∼ 0.5). 2 mL aliquots were collected by centrifugation at 1000 x *g* for 5 min. Cell pellets were resuspended in 450 μl 1X PBS (pH 7.4) and 50 μl of each SHAPE compound were added, followed by brief vortexing. Cells were allowed to react at 37°C for 20 min, with moderate shaking, after which 500 μl of 1 M DTT were added, to quench the reaction. Cells were pelleted by centrifugation at 17,000 x *g* for 2 min (4°C) and supernatant discarded. Cell pellets were then resuspended in 100 μl of Buffer A [10 mM Tris pH 8.0; 100 mM NaCl], supplemented with 5 U SUPERase•In™ RNase Inhibitor, by vigorously vortexing for 30 sec. 25 μl of Buffer B [50 mM EDTA; 120 mM Tris pH 8.0], supplemented with 100 μg/mL final lysozyme, were then added, followed by 30 sec vortexing. Samples were incubated 1 min at room temperature, after which 125 μl of Buffer C [0.5% Tween-20; 0.4% Sodium deoxycholate; 2 M NaCl; 10 mM EDTA] were added to lyse protoplasts. Samples were then incubated at room temperature for 5 min, after which 100 μl of lysate were transferred to 1 mL ice-cold QIAzol and thoroughly mixed by vortexing.

### *In vivo* probing of *B. subtilis* cells

A single colony of *Bacillus subtilis* 168 was picked and inoculated in LB medium without antibiotics, then grown to exponential phase (OD_600_ ∼ 0.5). 2 mL aliquots were collected by centrifugation at 1000 x *g* for 5 min. Cell pellets were resuspended in 450 μl 1X PBS (pH 7.4) and 50 μl of each SHAPE compound were added, followed by brief vortexing. Cells were allowed to react at 37°C for 20 min, with moderate shaking, after which 500 μl of 1 M DTT were added, to quench the reaction. Cells were pelleted by centrifugation at 17,000 x *g* for 2 min (4°C) and supernatant discarded. Cell pellets were then resuspended in 200 μl TE Buffer [10 mM Tris pH 8.0; 1 mM EDTA pH 8.0], supplemented with 1.1% final SDS. 0.25 g glass beads were then added and samples were beated in a Mini-Beadbeater-24 (Glen Mills) for 1 min. Samples were then incubated on ice for 1 min, after which beating was repeated once. 1 mL ice-cold QIAzol was then added, followed by vigorous vortexing. Samples were then incubated at 70°C for 20 min, to allow complete cell lysis.

### *In vivo* probing of HEK293 cells

HEK293 cells were grown in DMEM (4.5 g/L D-Glucose), supplemented with 10% heat-inactivated FBS, 0.1 mM NEAA, 1 mM Sodium Pyruvate, 25 U/mL penicillin and 25 μg/mL streptomycin, at 37°C (5% CO_2_), to ∼75% confluence. Cells were then covered with a thin layer of Trypsin-EDTA (0.25%) solution, incubated at room temperature for 1 min, then dissociated by pipetting up and down with complete medium. Cells were centrifuged at 180 x *g* for 1 min and medium discarded. Cells were then resuspended in 450 μl 1X PBS (pH 7.4) and 50 μl of each SHAPE compound were added, followed by gentle mixing by tapping the tube. Cells were allowed to react at 37°C for 15 min, with moderate shaking, after which 500 μl of 1 M DTT were added, to quench the reaction. Cells were pelleted by centrifugation at 10,000 x *g* for 1 min (4°C) and supernatant discarded. Cells pellets were lysed by direct addition of 1 mL ice-cold QIAzol, followed by a brief incubation at 56°C for 5 min to completely dissolve cell aggregates.

### Total RNA extraction

After samples have been collected in 1 mL QIAzol, 200 μl of chloroform were added, followed by vigorous vortexing for 15 sec. Samples were then incubated at room temperature for 2 min, after which they were centrifuged at 12,500 x *g* for 15 min (4°C). After centrifugation, the upper aqueous phase was transferred to a clean 2 mL tube, and mixed with 2 volumes of 100% ethanol by vigorous vortexing. The entire volume was then transferred to an RNA Clean & Concentrator™-5 column (Zymo Research, cat. R1015) and RNA was purified as per manufacturer instructions. RNA integrity was ensured by gel electrophoresis on a denaturing 2% agarose gel. Besides samples treated with I6, all other samples appeared to be perfectly intact. Before proceeding to library preparation, traces of genomic DNA were removed by treatment with 2 U TURBO™ DNase (ThermoFisher Scientific, cat. AM2239) at 37°C for 20 min.

### SHAPE-MaP library preparation

Total RNA was first fragmented to a median size of 150 nt by incubation at 94°C for 8 min in RNA Fragmentation Buffer [65 mM Tris-HCl pH 8.0; 95 mM KCl; 4 mM MgCl_2_], then purified with NucleoMag NGS Clean-up and Size Select beads (Macherey Nagel, cat. 744970), supplemented with 10 U SUPERase•In™ RNase Inhibitor, and eluted in 2 μl (for SSII and HIV RT) or 2.5 μl (for TGIRT-III) NF H_2_O. Eluted RNA was supplemented with 0.5 μl 20 μM random hexamers and either 0.25 μl (for SSII) or 0.5 μl (for HIV RT and TGIRT-III) dNTPs (10 mM each), then incubated at 70°C for 5 min and immediately transferred to ice for 1 min. Reverse transcription reactions were conducted in a final volume of 5 μl. For HIV RT, reaction was supplemented with 1 μl 5X RT Buffer [250 mM Tris-HCl pH 8.3; 375 mM KCl; 40 mM MgCl_2_], 0.5 μl DTT 0.1 M, 5 U SUPERase•In™ RNase Inhibitor and 5 U HIV RT (Worthington Biochemical Corporation, cat. HIVRT). For TGIRT-III, reaction was supplemented with 1 μl 5X RT Buffer [250 mM Tris-HCl pH 8.3; 375 mM KCl; 15 mM MgCl_2_], 0.25 μl DTT 0.1 M, 5 U SUPERase•In™ RNase Inhibitor and 50 U TGIRT-III RT (InGex, cat. TGIRT-50). For SSII, reaction was supplemented with 1 μl 5X RT Buffer [250 mM Tris-HCl pH 8.3; 375 mM KCl], 0.5 μl DTT 0.1 M, 0.25 μl MnCl_2_ 120 mM, 5 U SUPERase•In™ RNase Inhibitor and 50 U SuperScript II RT (ThermoFisher Scientific, cat. 18064014). Reactions were incubated at 25°C for 10 min to allow partial primer extension, followed by 2 h at 37°C (for HIV RT), 2 h at 57°C (for TGIRT-III), or 3 h at 42°C (for SSII). SSII and HIV RT were heat-inactivated by incubating at 75°C for 20 min. As TGIRT-III tightly binds the cDNA-RNA complex, 0.5 μg of Proteinase K were added and reaction was incubated at 37°C for 20 min, after which 0.5 μl of a 1:2 dilution of protease inhibitor cocktail (Sigma Aldrich, cat. P9599) in water was added to stop the reaction. For SSII, the buffer exchange step was omitted. Instead, 6 mM final EDTA was added to chelate Mn^2+^ ions, followed by 5 min incubation at room temperature and addition of 6 mM final MgCl_2_. Reverse transcription reactions were then used as input for the NEBNext® Ultra II Non-Directional RNA Second Strand Synthesis Module (New England Biolabs, cat. E6111L). Second strand synthesis was performed by incubating 1 h at 16°C, as per manufacturer instructions. DsDNA was purified using NucleoMag NGS Clean-up and Size Select beads, and used as input for the NEBNext® Ultra™ II DNA Library Prep Kit for Illumina® (New England Biolabs, cat. E7645L), following manufacturer instructions.

### Analysis of SHAPE-MaP data

All the relevant data analysis steps have been conducted using RNA Framework v2.6.9 (28). Reads pre-processing and mapping was performed using the *rf-map* module (parameters: *-ctn -cmn 0 -cqo -cq5 20 -b2 -mp “--very-sensitive-local”*). Reads were trimmed of terminal Ns and low-quality bases (Phred < 20). Reads with internal Ns were discarded. Mapping was performed using the “very-sensitive-local” preset of Bowtie2 (29). The mutational signal was derived using the *rf-count* module (parameters: *-m -rd*), enabling the right re-alignment of deletions. Grid search of optimal *slope*/*intercept* pairs was performed using the *rf-jackknife* module (parameters: *-rp “-md 600 -nlp” -x*) and ViennaRNA package v2.4.11 (30), disallowing isolated base-pairs, setting the maximum base-pairing distance to 600 nt and enabling the relaxed structure comparison mode (9) (a basepair *i/j* is considered to be correctly predicted if any of the following pairs exist in the reference structure: *i/j*; *i-1/j*; *i*+*1/j*; *i/j-1*; *i/j*+*1*).

### Reference structure of rRNAs

16S and 23S rRNA reference structures for *E. coli* and *B. subtilis* were obtained from the Comparative RNA Web (31). 18S and 28S rRNA reference structures for *H. sapiens* were obtained from RNAcentral (32).

### Generation of Receiver Operating Characteristic (ROC) curves

For the generation of ROC curves, the mutation frequency threshold was linearly varied from 0 to 1 with a step of 0.001, calculating at each increment the number of unpaired (true positive) and base-paired (true negative) residues (according to the accepted reference rRNA structures), whose mutation frequency exceeded that of the threshold. Terminal base-pairs (beginning/end of helices and isolated base-pairs) were excluded. For *in vivo* samples, only solvent-accessible bases were considered, as previously described (33). For solvent accessible surface area (SASA) calculation, ribosome structures were obtained from the Protein Data Bank, using accessions 5IT8 (*E. coli*), 6HA1 (*B. subtilis*) and 4UG0 (*H. sapiens*) in CIF format and converted to PDB format using PyMOL (v2.3.5). SASA per residue was then calculated using POPS (v3.1.2) and a probe radius of 3 Å. Accessible residues were defined as those residues with a SASA greater than 2 Å. Area under the curve (AUC) was calculated using the MESS package (v0.5.6) for R.

### Signal-to-noise estimation

From reference rRNA structures in dot-bracket notation, we extracted all hairpin-loops with a stem length ≥ 3 and a loop size ≥ 3, using the regular expression “([\(]{3}\.{3,}[\)]{3})”. For each hairpin-loop we then calculated the median reactivity on the first and last 2 bases of the loop and the median reactivity on the 3 bases of the stem preceding and following the loop. The signal-to-noise value was then calculated as the ratio between the median reactivity across all the loops and the median reactivity across all the stems.

## RESULTS

### Comparison of RT enzymes for SHAPE-MaP

In a first attempt to optimize the signal-to-background of SHAPE mutational profiling approaches (SHAPE-MaP), we sought to find the best reverse transcription conditions. To this end, we extracted deproteinized total RNA from *Escherichia coli* cells, under conditions previously shown to preserve the native RNA folding (9, 24, 34) and probed it in solution with NAI (or neat DMSO as a control). We then performed reverse transcription using three different reverse transcriptases: SuperScript II (SSII) in Mn^2+^-containing buffer, thermostable group II intron RT (TGIRT-III) and HIV RT (23, 25, 35). These enzymes have been previously shown to be able to generate mutational signatures in cDNA when reading through SHAPE adducts (SSII), DMS-induced alkylations (SSII and TGIRT-III), or N-cyclohexyl-N’-(2-morpholinoethyl)carbodiimide metho-p-toluenesulfonate (CMCT) adducts (HIV RT). Analysis of the distribution of per-base mutation frequencies across 16S and 23S rRNAs showed that, while TGIRT-III has lower background mutation frequencies compared to SSII (median DMSO control: 0.75 × 10^−3^ for TGIRT-III vs. 2.19 × 10^−3^ for SSII) as previously reported (23), SSII shows a higher and more significant increase in the mutation rates measured in the NAI treated sample with respect to the DMSO control (ratio median NAI vs. DMSO: 2.95 for SSII, 1.37 for TGIRT-III and 1.16 for HIV RT; Figure S1A). Accordingly, Receiver Operating Characteristic (ROC) curves built with respect to unpaired versus paired residues from phylogenetically-inferred *E. coli* rRNA secondary structure models (31) and inspection of SHAPE-MaP signals confirmed the higher ability of SSII to translate SHAPE adducts on structurally-flexible bases into a mutational signature (Figure S1B-C). Altogether, these results identify SSII in Mn^2+^-containing buffer as the optimal RT enzyme for SHAPE-MaP experiments.

### Synthesis of six novel candidate SHAPE reagents

While designing new candidate SHAPE compounds, we sought to meet the following requirements: 1) permeable to biological membranes (suitable for *in vivo* probing experiments); 2) high signal over background (deriving from a good trade between slow hydrolysis in water, reactivity with RNA and membrane permeability); 3) easy and safe to synthesize. The two best-characterized *in vivo* SHAPE reagents to date are NAI and FAI. These compounds can be easily synthesized via the Carbonyldiimidazole (CDI)-mediated activation of carboxylic acids, resulting in the corresponding carboxylic acid imidazolide, an activated carbonyl, that can acylate nucleophiles present in the system, especially 2’ hydroxyl groups of RNA. We decided to adopt the same strategy, trying to optimize the acylating group. To this end, we selected seven commercially-available carboxylic acids, preferring those containing nitrogen groups or for which the putative adduct-forming group had a pKa close to (or slightly above) 7 (Table 1; see Materials and Methods).

**Table 1.**
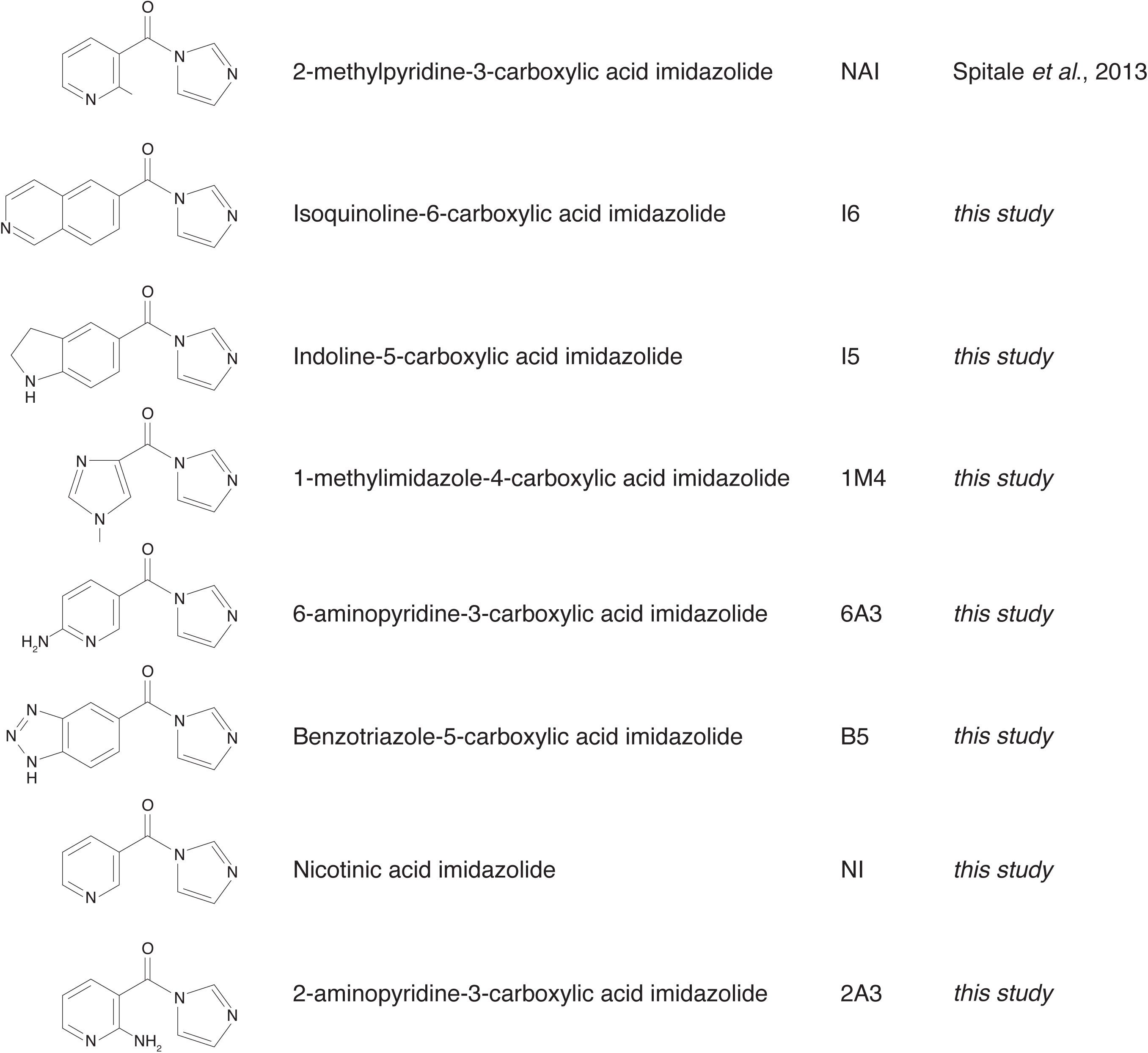
SHAPE reagents used in this study. Chemical structure, full name and acronym of the SHAPE reagents tested in this study. NAI, used as a standard, is reported for comparison.

|  |  |  |  |
| --- | --- | --- | --- |
|  | 2-methylpyridine-3-carboxylic acid imidazolide | NAI | Spitale *et al.*, 2013 |
|  | Isoquinoline-6-carboxylic acid imidazolide | I6 | *this study* |
|  | Indoline-5-carboxylic acid imidazolide | I5 | *this study* |
|  | 1-methylimidazole-4-carboxylic acid imidazolide | 1M4 | *this study* |
|  | 6-aminopyridine-3-carboxylic acid imidazolide | 6A3 | *this study* |
|  | Benzotriazole-5-carboxylic acid imidazolide | B5 | *this study* |
|  | Nicotinic acid imidazolide | NI | *this study* |
|  | 2-aminopyridine-3-carboxylic acid imidazolide | 2A3 | *this study* |

We then evaluated all reagents for their ability to react with structurally-flexible RNA residues using *E. coli ex vivo* deproteinized total RNA and NAI probing as the gold standard. We further discarded isoquinoline-6-carboxylic acid imidazolide (I6I) as it was too reactive, hence resulting in very fast RNA degradation (data not shown). When looking at overall distributions of mutation rates, indoline-5-carboxylic acid imidazolide (I5) and 1-methylimidazole-4-carboxylic acid imidazolide (1M4) showed a lower reactivity towards RNA compared to NAI (median: 3.07 × 10^−3^ for I5, 4.50 × 10^−3^ for 1M4). Oppositely, the other four compounds, nicotinic acid imidazolide (NIC), benzotriazole-5-carboxylic acid imidazolide (B5), 6-aminopyridine-3-carboxylic acid imidazolide (6A3) and 2-aminopyridine-3-carboxylic acid imidazolide (2A3), all showed increased reactivity compared to NAI (median: 8.90 × 10^−3^ for NIC, 9.90 × 10^−3^ for B5, 11.9 × 10^−3^ for 6A3 and 15.2 × 10^−3^ for 2A3), with 2A3 having a ∼2.4X higher mutation frequency than NAI (Figure 1A). ROC curves built with respect to unpaired versus paired residues from *E. coli* rRNAs revealed nearly comparable accuracies for 6A3, B5, NIC and 2A3 compared to NAI, with NIC and 2A3 showing a slightly higher area under the curve (AUC), independently of DMSO background signal subtraction (Figure 1B and S2A). This intrinsically implies that these two reagents have a slightly higher signal-to-noise ratio with respect to NAI. To systematically quantify the actual signal-to-noise ratio of the different reagents, we extracted all the individual hairpin-loops from the known structures of *E. coli* 16S and 23S rRNAs, and calculated the median mutation frequency across the first and last 2 bases of the loop, versus the 3 stem bases preceding or following the loop (Figure S2B). Notably, the ratio between the median loop reactivity and the median stem reactivity for 2A3 (4.01), B5 (4.01) and NIC (4.19) was slightly higher than that of NAI (3.59), confirming that these reagents might represent better choices for *in vivo* RNA probing.

**Figure 1.**
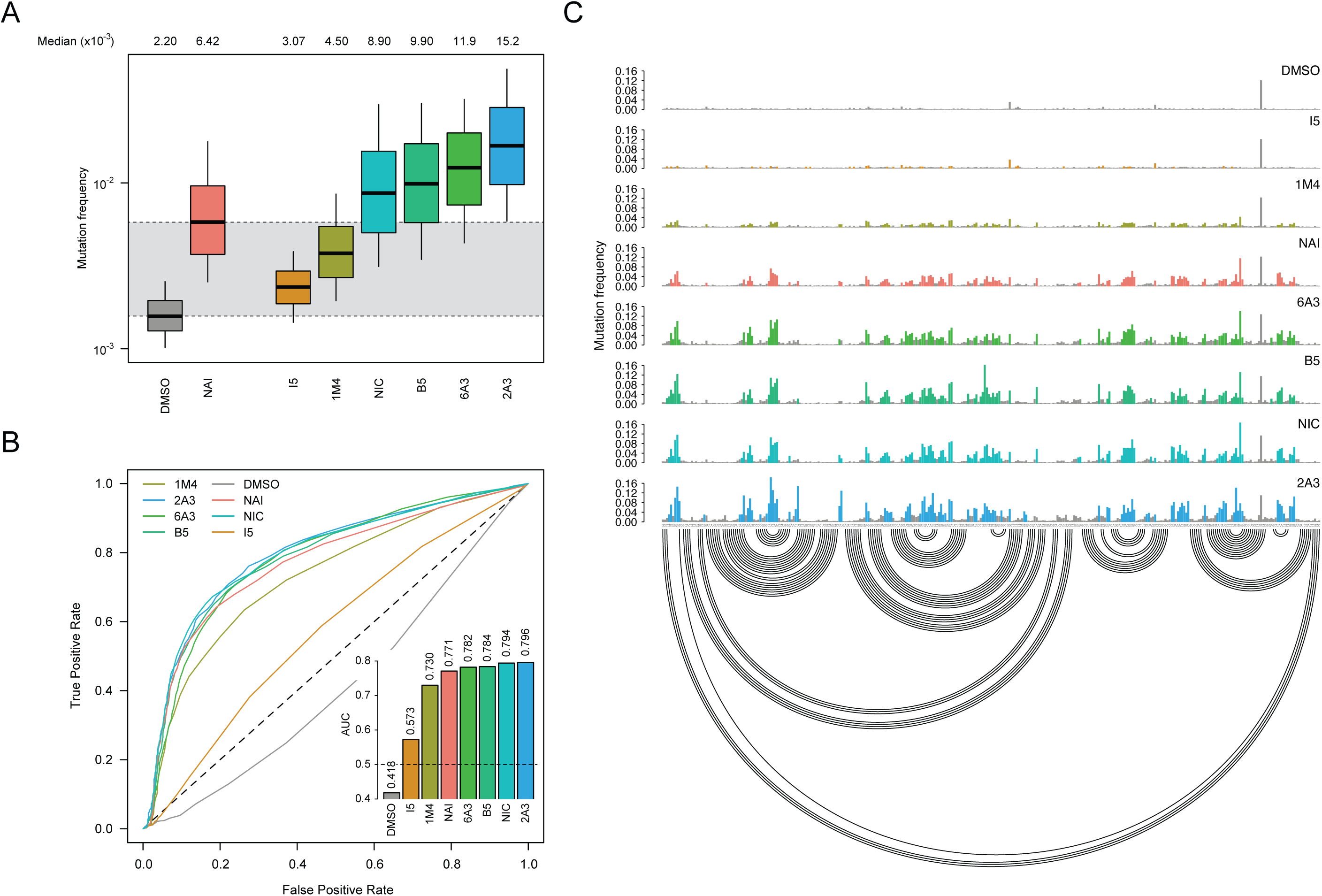
Comparison of SHAPE reagents under *ex vivo* conditions. (A) Boxplot of SHAPE−MaP mutation frequencies for *E. coli* 16S and 23S rRNAs probed *ex vivo* after deproteinization. Box plots span the interquartile range (from Q1 to Q3). The grey area spans from the median in the DMSO sample (control) to the median in the NAI sample (reference). (B) ROC curve for all tested SHAPE reagents, calculated with respect to the accepted phylogenetically-inferred 16S and 23S rRNA structures from CRW. The inset reports the area under the curve (AUC) for each compound. (C) Sample of SHAPE−MaP mutation frequencies for all tested compounds across a region spanning nucleotides 567 to 884 of *E. coli* 16S rRNA. Colored bases are those whose mutation frequencies exceed by 2-folds the median mutation frequency in the analyzed region. The accepted structure is reported as an arc plot.

### 2A3 outperforms NAI at probing RNA *in vivo*

Spurred by these promising results, we decided to test the ability of these newly synthesized compounds to permeate biological membranes to probe RNA within living cells. To this end, we employed *E. coli, B. subtilis* and HEK293 cells as representatives of the three major cell types in research labs: Gram-negative, Gram-positive and mammalian cells.

At first, we looked again into the distribution of mutation frequencies for the different reagents. Unexpectedly, while the observed distributions for HEK293 cells approximately mirrored those observed on *ex vivo*-modified RNAs (Figure S3A), we observed that, for both *E. coli* and *B. subtilis*, mutation frequency distributions for most reagents, including NAI, were extremely close to those of the control DMSO sample (Figure 2A and S3B). On the other hand, 2A3 showed significantly higher mutation frequencies compared to NAI (ratio median 2A3 vs. NAI: 2.74 in *E. coli*, 1.61 in *B. subtilis* and 1.77 in HEK293), independently of the probed cell type, partially followed by B5 in *B. subtilis* and NIC *in* HEK293 cells.

**Figure 2.**
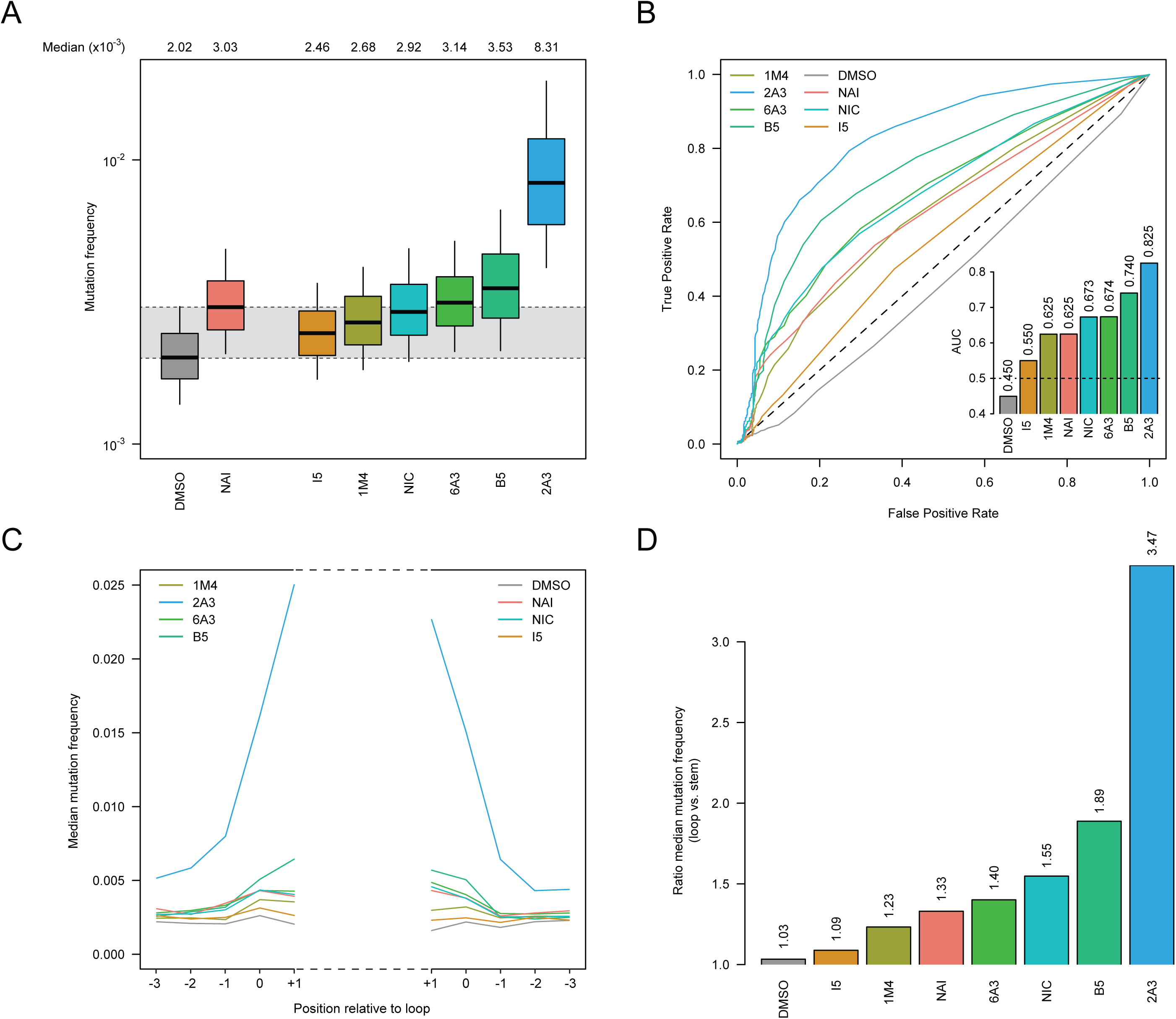
Comparison of SHAPE reagents under *in vivo* conditions. (A) Boxplot of SHAPE−MaP mutation frequencies for *E. coli* 16S and 23S rRNAs probed *in vivo*. Box plots span the interquartile range (from Q1 to Q3). The grey area spans from the median in the DMSO sample (control) to the median in the NAI sample (reference). (B) ROC curve for all tested SHAPE reagents, calculated on solvent-exposed residues in the crystal structure of the *E. coli* ribosome (PDB: 5IT8), with respect to the accepted phylogenetically-inferred 16S and 23S rRNA structures from the Comparative RNA Web (31). The inset reports the AUC for each compound. (C) Median *in vivo* SHAPE-MaP mutation frequencies across all stem-loops in the accepted *E. coli* 16S and 23S rRNA structures. Bases are numbered relatively to the loop. Positions −3 to −1 correspond to stem bases, while positions 0 and +1 correspond to loop bases. (D) Ratio between the median loop mutation frequency and the median stem mutation frequency, calculated on all stem-loops from C.

Oppositely to what we observed for *ex vivo-*probed RNAs, ROC curves built with respect to unpaired versus paired residues from *E. coli, B. subtilis* and *H. sapiens* rRNAs (calculated on residues that are solvent-exposed in the respective ribosome crystal structures; see Materials and Methods) revealed a marked difference in sensitivity/specificity of NAI compared to 2A3, with 2A3 showing up to 17-20% increased discrimination of unpaired versus paired residues, independently of DMSO background signal subtraction (AUC *E. coli*: 0.825/0.874 for 2A3 -/+ DMSO subtraction, 0.625/0.708 for NAI -/+ DMSO subtraction; Figure 2B and S4-5). Accordingly, quantitation of SHAPE-MaP signal on loop versus stem regions confirmed the strikingly higher signal-to-noise ratio of 2A3 (*E. coli*: 3.47; *B. subtilis*: 2.36; HEK293: 3.01) with respect to NAI (*E. coli*: 1.33; *B. subtilis*: 1.56; HEK293: 2.08), independently of the examined cell type (Figure 2C-D and S6).

Deeper inspection of SHAPE-MaP signals revealed that, regions of bacterial ribosomes that were almost completely blind to NAI and most reagents, were instead partially accessible to B5 and readily probed with high efficiency by 2A3, with mutational signatures in strong agreement with the known rRNA structure (Figure 3A and S7). In these regions, not only 2A3 effectively probed most RNA loops, but also it better captured local nucleotide dynamics. As an example, base U_1032_ of *E. coli* 23S rRNA is hyperreactive to 2A3, but almost completely unreactive to any other tested reagent, including NAI (Figure 3A). Coherently, analysis of this base in the context of the crystal structure of the fully assembled *E. coli* ribosome (36) reveals its extreme flexibility, as it is twisted, completely flipping out of the parent helix (Figure 3B). Oppositely, a large loop of 19 contiguous single-stranded residues between positions 1124 and 1181 showed very low reactivities. Notably, this loop has a very low B factor in the crystal, suggesting that these residues are weakly dynamic (lowly flexible) in the context of the ribosome. In agreement with this observation, these residues became highly reactive towards 2A3 upon deproteinization (Figure S8). Similarly, the hyperreactivity of U1032 was abrogated upon rRNA deproteinization. In HEK293, although NAI readily accessed most regions of the ribosome, showing very similar reactivity patterns to the other tested reagents, we observed that probing with 2A3 (as well as with NIC) resulted in a stronger and neater SHAPE-MaP signal (Figure S9).

**Figure 3.**
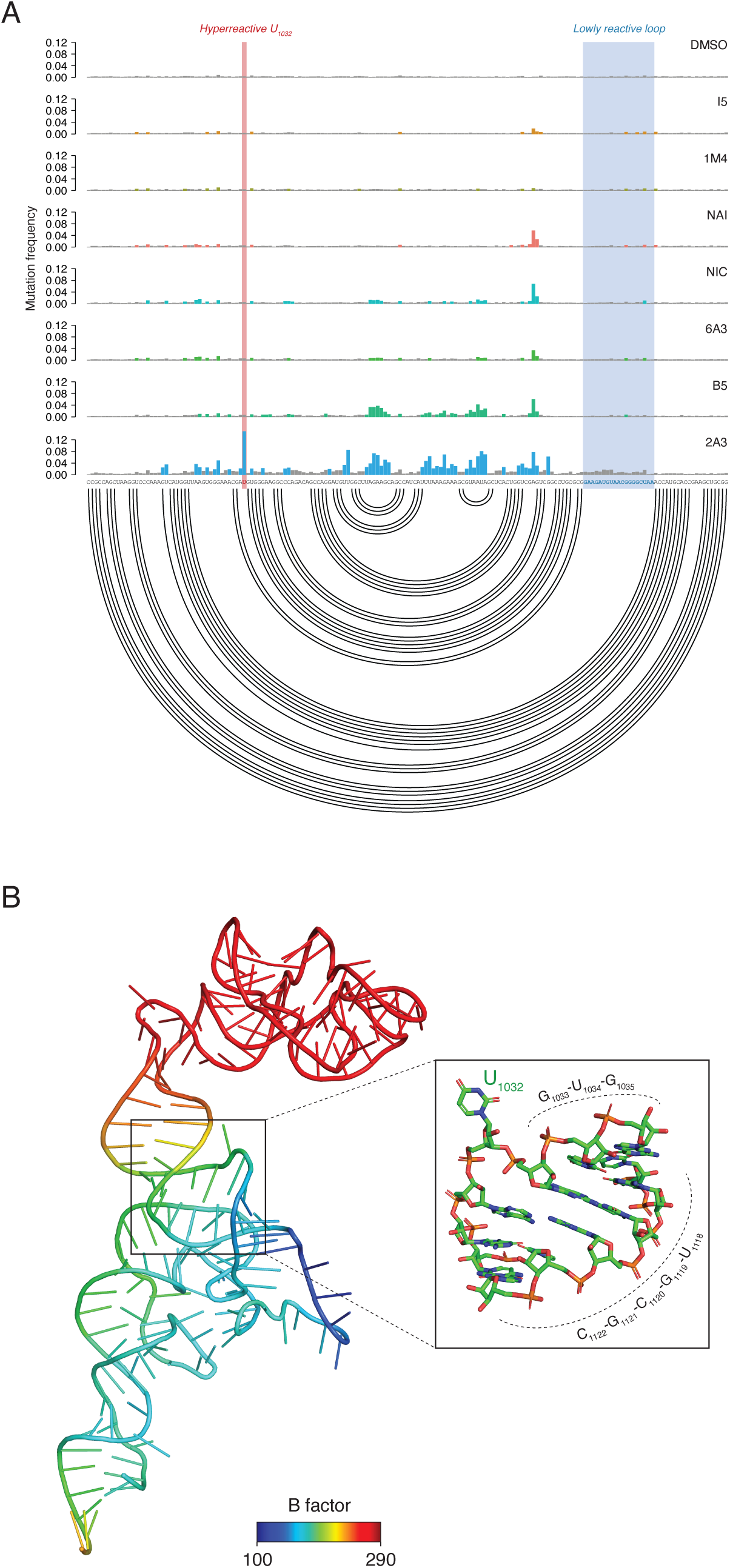
2A3 successfully queries regions of the ribosome that are blind to other reagents. (A) Sample of SHAPE−MaP mutation frequencies for all tested compounds across a region spanning nucleotides 990 to 1162 of *E. coli* 23S rRNA. Colored bases are those whose mutation frequencies exceed by 2-folds the median mutation frequency in the analyzed region. The accepted structure is reported as an arc plot. The hyperreactive residue U_1032_ and the unreactive stretch of 19 nucleotides are respectively marked in red and blue. (B) Three-dimensional model of the domain depicted in A, colored by B factor (PDB: 5IT8). The inset zooms on the helical region containing the 2A3-hyperreactive residue U_1032_.

### 2A3 markedly improves the accuracy of RNA structure prediction

To definitively prove the better suitability of 2A3 for *in vivo* RNA folding studies, we used SHAPE-inferred reactivities to perform experimentally-driven RNA structure modeling. SHAPE reactivities can be incorporated into thermodynamics-based prediction algorithms by converting them into pseudo-free energy contributions (9). This conversion relies on the use of two empirically-determined parameters, the *slope* (*m*) and the *intercept* (*b*). As these parameters depend on the experimental setup and on the employed reagent, we first used the ViennaRNA Package 2.0 (30) to perform a grid search of the optimal *m*/*b* pairs for NAI and 2A3 for *E. coli* rRNAs, in order to avoid skewing prediction results because of the use of the same *m*/*b* pair for both reagents (Figure 4A). For each *m*/*b* pair, we calculated the geometric mean of the prediction’s sensitivity (the fraction of known base pairs correctly predicted) and positive predictive value (PPV; the fraction of predicted pairs that are present in the known reference structure) for both 16S and 23S rRNAs. Notably, grid search already revealed that the accuracy of 2A3-driven predictions was largely independent of the employed slope/intercept pair and that, even the most accurate NAI-driven prediction (*m*: 2.2; *b*: −0.8; geometric mean of PPV/sensitivity: 0.67) was ∼10% less accurate than most 2A3-driven predictions, independently of the used *m*/*b* pair and ∼18% less accurate that the top-scoring 2A3-driven prediction (*m*: 1.0; *b*: −0.4; geometric mean of PPV/sensitivity: 0.85). As an example, the NAI-guided prediction resulted in the misfolding of the 3’ domain of the 16S rRNA (PPV: 0.65; sensitivity: 0.66), while the 2A3-guided prediction correctly recapitulated most of the known base-pairs from the accepted reference structure (PPV: 0.82; sensitivity: 0.90). A similar improvement was observed also when modeling the structure of *B. subtilis* rRNAs (Figure S10). Use of RNAstructure (37) rather than ViennaRNA produced similar results (Figure S11), hence demonstrating that our conclusions are software-independent.

**Figure 4.**
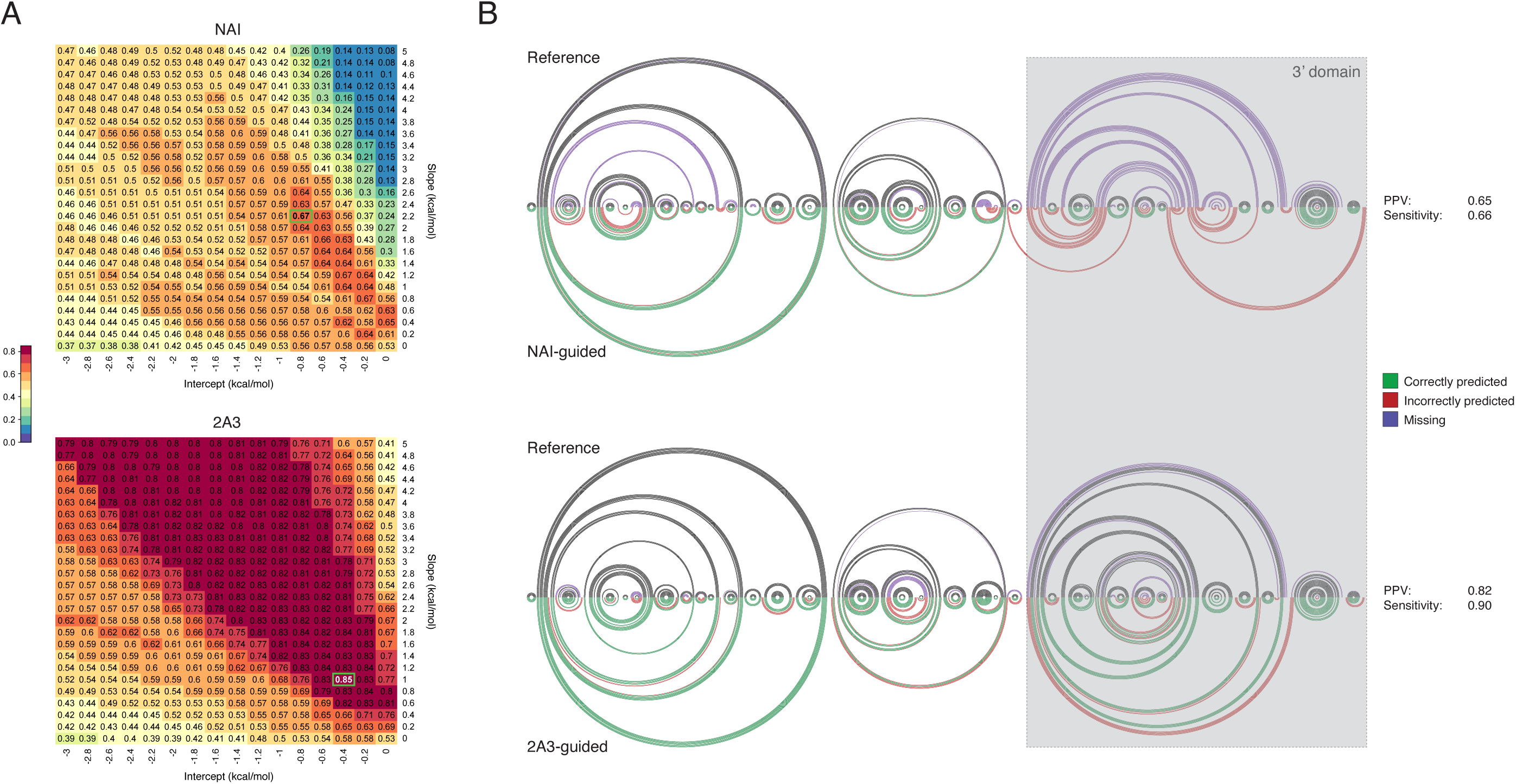
2A3 outperforms NAI at experimentally-driven RNA structure modeling. (A) Grid search (jackknifing) of optimal slope/intercept value pairs for *E. coli* 16S and 23S rRNAs *in vivo* probing data for NAI and 2A3. Values represent the geometric mean of sensitivity and PPV for the secondary structures predicted using each slope/intercept value pair. The chosen value pair is boxed in green. (B) Arc plot comparison of *E. coli* 16S rRNA reference structure (top), and structure inferred using either NAI-derived or 2A3-derived restraints (bottom). Black/green arcs correspond to correctly predicted base-pairs, violet arcs to non-predicted base-pairs and red arcs to mispredicted base-pairs. PPV and sensitivity for each prediction are indicated.

### Chemical characterization of 2A3

Analytical preparations of 2A3 in DMSO unexpectedly revealed that a secondary product was occurring. This reaction proceeded slowly at room temperature after the rapid initial formation of the desired imidazolide, 2A3. The resulting product was a cyclization reaction leading to the formation of 3-azaisatoic anhydride (3AIA). This unwanted side product increased with time, eventually reaching a maximum of approximately 90% as compared to 2A3 in solution during an overnight reaction in a sealed vessel. The formation of 2A3 proceeds rapidly as the 2-aminonicotinic acid after the addition of CDI. The conversion of 2A3 to 3AIA then proceeds at room temperature in the crude reaction mixture. The unwanted side reaction proceeds slowly and can be minimized with vigorous stirring under a constant flow of inert gas. To rule out the possibility that 3AIA rather than 2A3 was acting as an *in vivo* SHAPE probe, we obtained commercially-available pure 3AIA (Synchem, Germany) and used it to probe living *E. coli* cells. Notably, 3AIA showed modification frequencies indistinguishable from those of the DMSO control *in vivo* (Figure S12), hence confirming that 2A3 was indeed responsible for the previously observed RNA modification.

## DISCUSSION

We have introduced here 2-aminopyridine-3-carboxylic acid imidazolide (2A3; Figure 5), a novel powerful and accurate SHAPE reagent for the interrogation of RNA structures, both *in vitro* and *in vivo*. Briefly, after synthesizing six new candidate SHAPE reagents, we evaluated their ability to modify RNA in cell-free conditions. Although all compounds were able to react with RNA to different extents, four of them (B5, NIC, 6A3 and 2A3) showed partial improvements with respect to NAI. Surprisingly, when tested *in vivo*, 2A3 significantly outperformed NAI and all the other tested reagents under all the analyzed conditions, with the most striking differences being observed for bacterial cells. Indeed, for both *E. coli* and *B. subtilis*, several regions of the ribosome resulted to be blind to probing by NAI and most of the other tested reagents, while they were readily probed by 2A3. In this context, 2A3 better captured local nucleotide dynamics, as confirmed by in depth analysis of available ribosome 3D structures. This reveals that, besides showing a higher ability to react with RNA, 2A3 has also a greater capacity to permeate biological membranes, being able to diffuse through both the double membrane of Gram-negatives and the peptidoglycan of Gram-positives.

**Figure 5.**
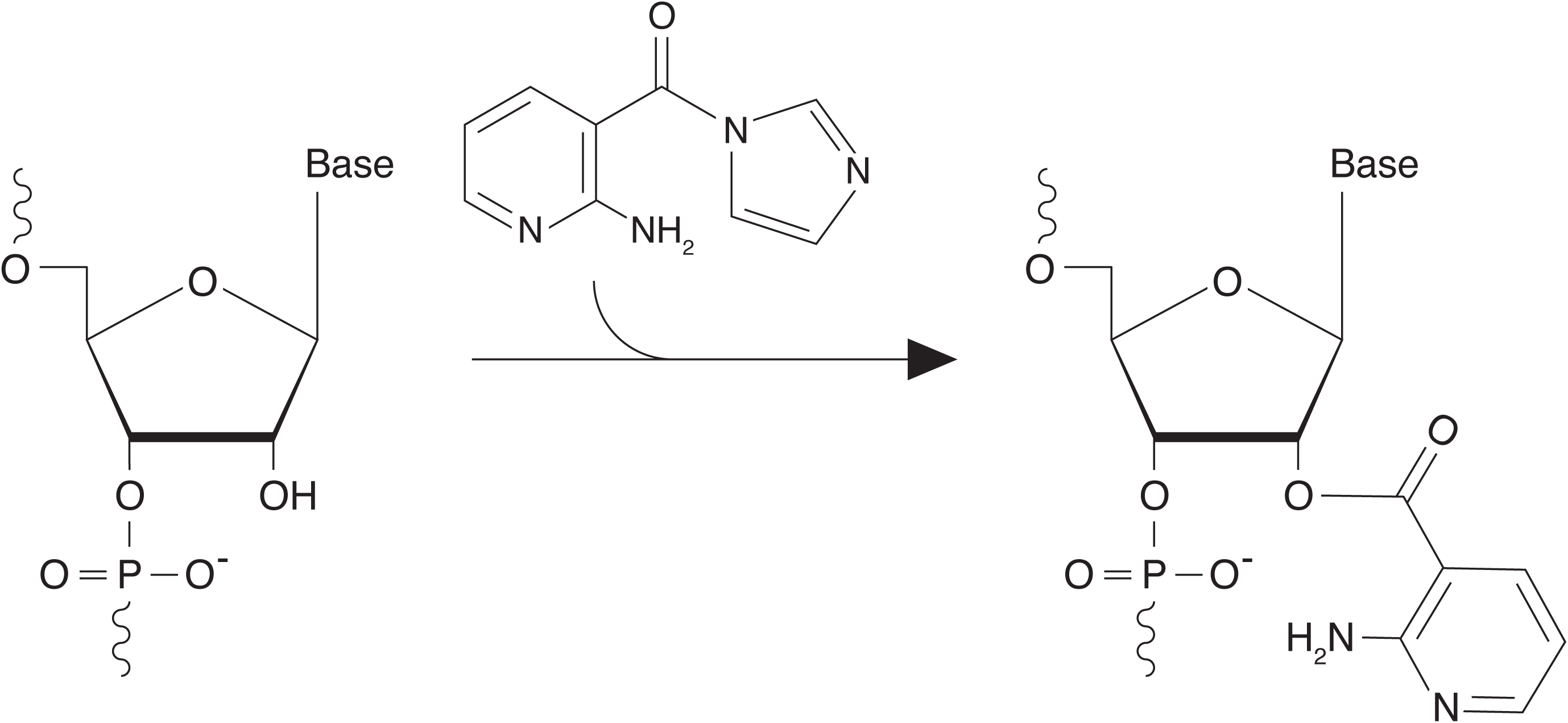
Mechanism of 2A3 reaction with RNA. Reaction of 2A3 with the 2’-OH of structurally-flexible RNA residues, resulting in the formation of a 2’-O-adduct.

Furthermore, we showed that, the higher signal-to-noise ratio achieved by 2A3 results in a significantly higher RNA structure prediction accuracy when performing 2A3-informed RNA folding. It is worth noticing that 2A3, as opposed to NAI, outperforms probing with dimethyl sulfate (DMS) as well. Indeed, probing with DMS usually shows a higher signal-to-noise ratio with respect to NAI, hence resulting in more accurate experimentally-informed RNA structure predictions (see for example Figure 3B in (28) and Figure 4A in this work; geometric mean of PPV/sensitivity on *E. coli* 16S and 23S rRNAs: 0.71 for DMS, 0.67 for NAI) even if providing information on roughly only a half of RNA bases.

Compounds NIC, 2A3 and 6A3 were designed around the nicotinic acid core of NAI, where 2A3 and 6A3 were chosen to be electronically similar but sterically different. The electronic effects predict similar reactivities for 2A3 and 6A3, but the amino group ortho to the activated carbonyl in 2A3 is expected to decrease the reactivity due to steric considerations much the way the ortho methyl in NAI decreases the reactivity of NAI relative to NIC. Surprisingly, 2A3 was found to be more reactive than 6A3 as shown by their hydrolysis half-lives (Table S1). It may be that 2A3 falls in a “sweet” spot for reactivity compared to the other tested electrophiles both *in vitro* and *in vivo*. In summary, 2A3 represents a concrete and significant advance in our ability to interrogate RNA structures in living cells. We can anticipate that, being able to readily probe even regions of ribonucleoprotein complexes that are normally hidden to other compounds, 2A3 will rapidly overtake conventional SHAPE reagents, revealing new and previously unobserved features of the *in vivo* RNA structurome.

## AVAILABILITY

SHAPE-MaP data has been deposited to the Gene Expression Omnibus (GEO) database, under the accession GSE154563. Data for *ex vivo* probing of *E. coli* RNA with DMSO and NAI has been previously deposited under the accession GSE151327, as part of another study. Additional processed files are available at: http://www.incarnatolab.com/datasets/2A3_Marinus_2020.php.

## ACKNOWLEDGMENTS

We would like to acknowledge Prof. Giovanni Maglia (University of Groningen) for critical discussion concerning the design of the new SHAPE compounds.

