## Supplementary Information for "A novel SHAPE reagent enables the analysis of RNA structure in living cells with unprecedented accuracy"

### for

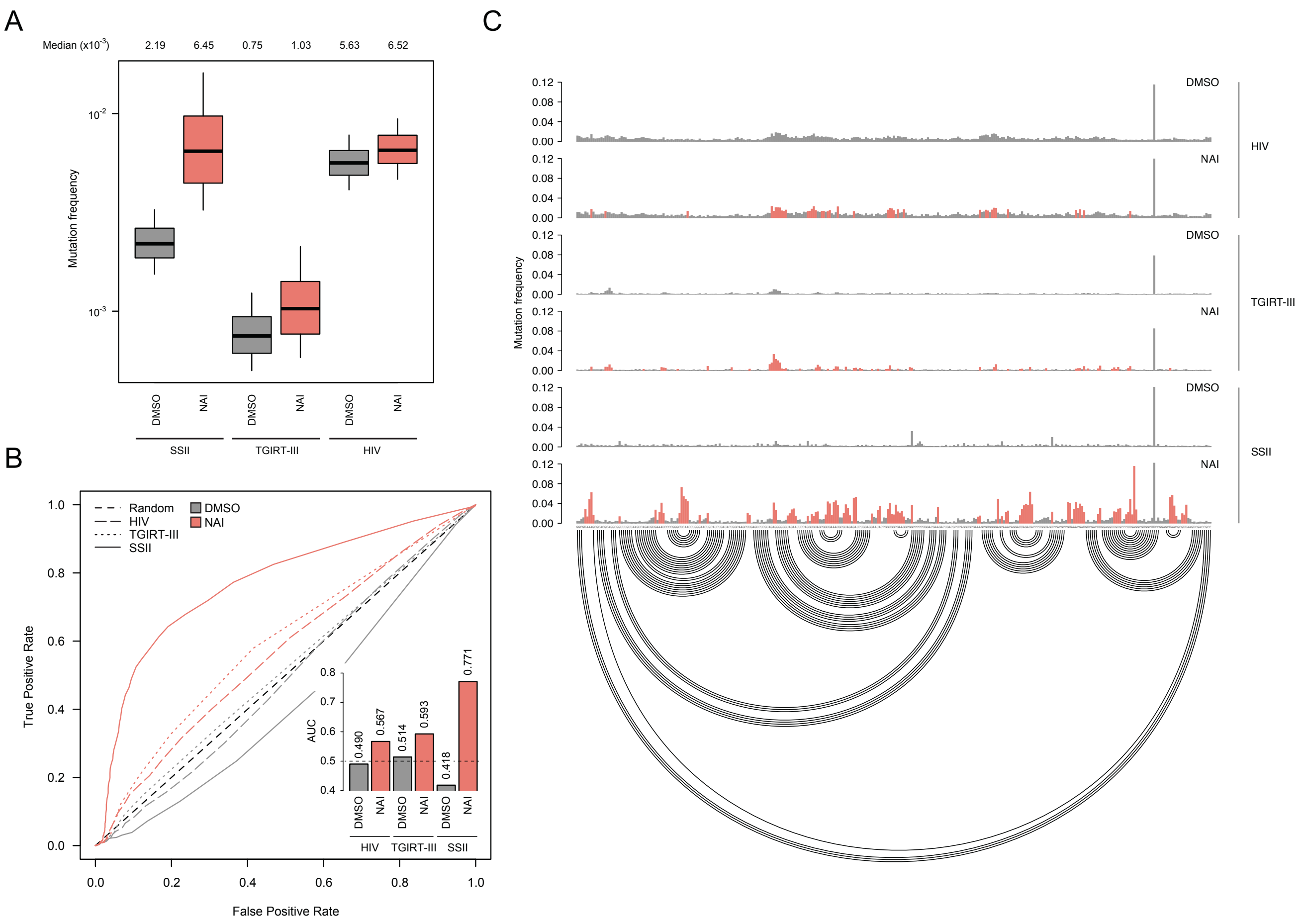


**Figure S1.** (A) Boxplot of SHAPE–MaP mutation frequencies for *E. coli* 16S and 23S rRNAs probed *ex vivo* after deproteinization. Box plots span the interquartile range (from Q1 to Q3). (B) ROC curve for all tested reverse transcriptases, calculated with respect to the accepted phylogenetically-inferred 16S and 23S rRNA structures from CRW. The inset reports the area under the curve (AUC) for each condition. (C) Sample of SHAPE–MaP mutation frequencies for all tested conditions across a region spanning nucleotides 567 to 884 of *E. coli* 16S rRNA. Colored bases are those whose mutation frequencies exceed by 2-folds the median mutation frequency in the analyzed region. The accepted structure is reported as an arc plot.

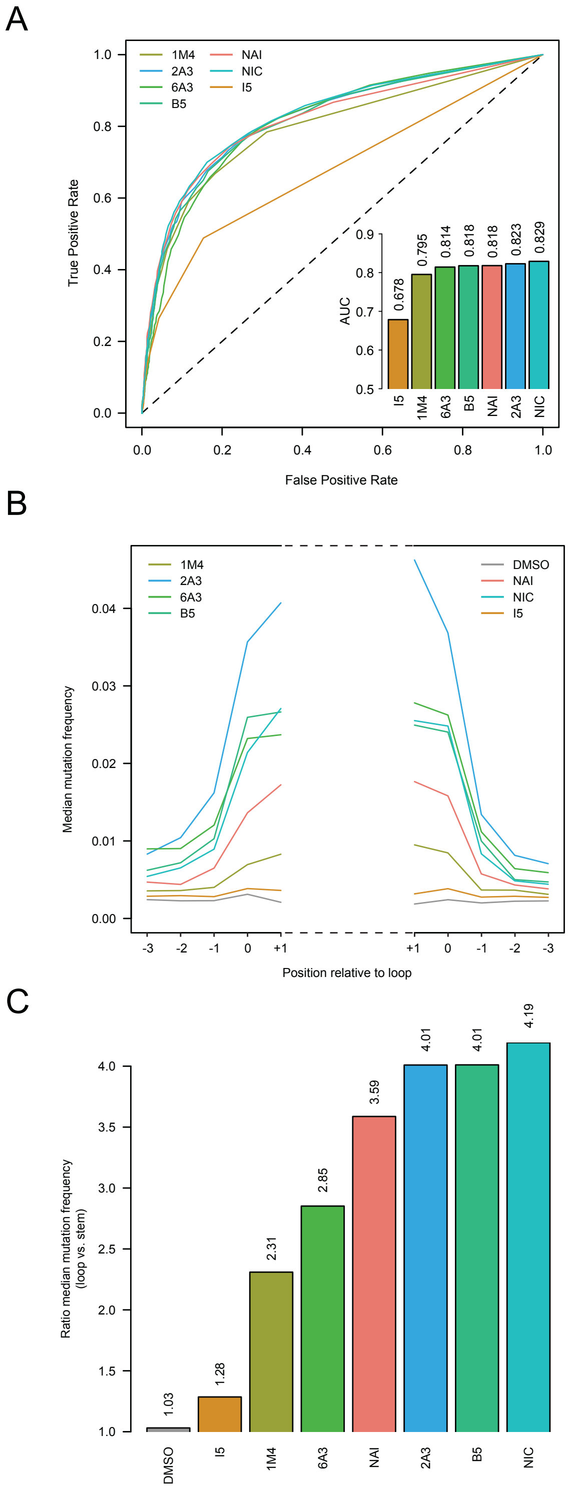


**Figure S2.** (A) ROC curve for all tested SHAPE reagents, calculated with respect to the accepted phylogenetically-inferred *E. coli* 16S and 23S rRNA structures from CRW. The inset reports the AUC for each compound. (B) Median SHAPE-MaP mutation frequencies across all stem-loops in the accepted 16S and 23S rRNA structures. Bases are numbered relatively to the loop. Positions -3 to -1 correspond to stem bases, while positions 0 and +1 correspond to loop bases. (C) Ratio between the median loop mutation frequency and the median stem mutation frequency, calculated on all stem-loops from B.

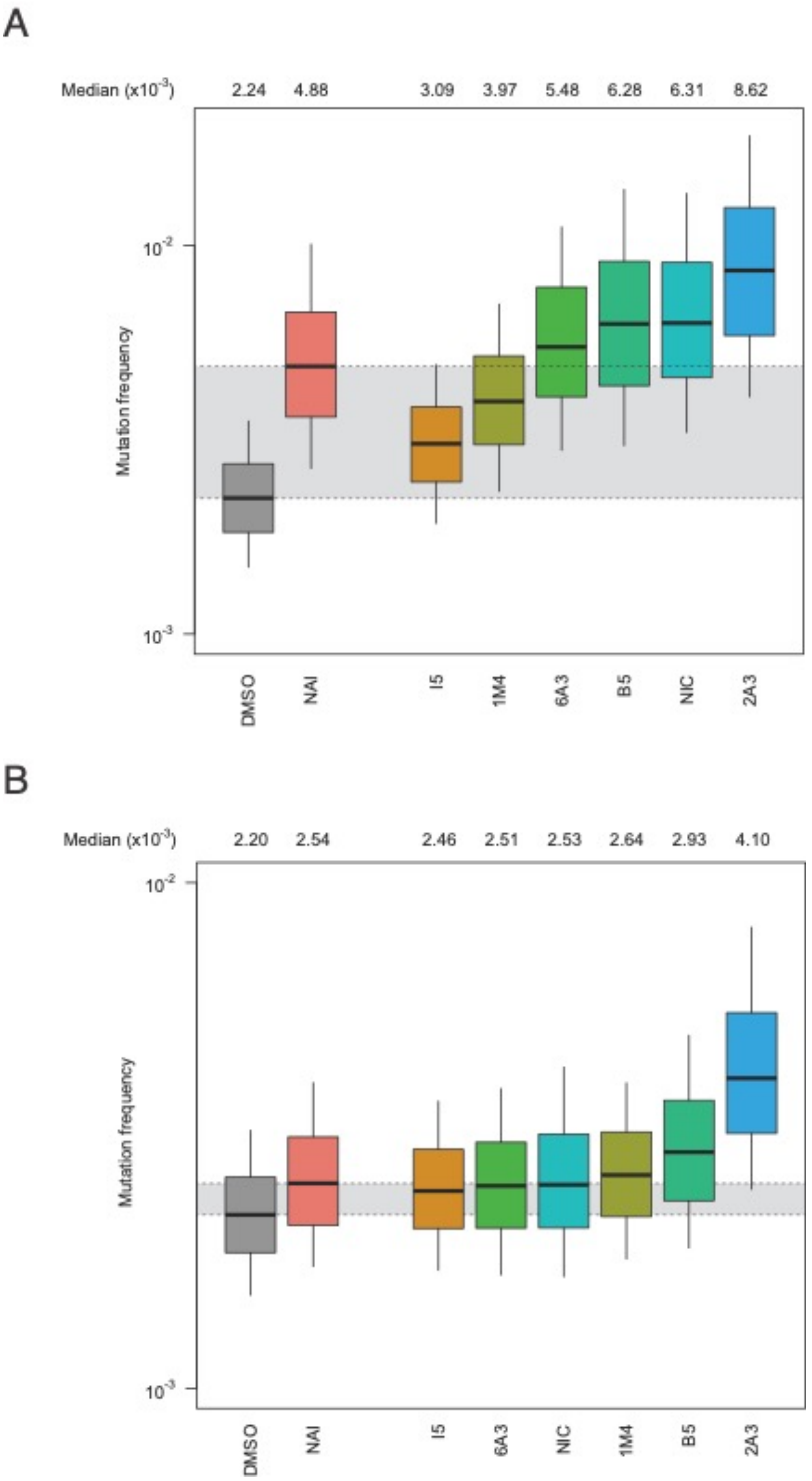


**Figure S3.** (A) Boxplot of SHAPE–MaP mutation frequencies for *H. sapiens* 18S and 28S rRNAs probed *in vivo*. Box plots span the interquartile range (from Q1 to Q3). The grey area spans from the median in the DMSO sample (control) to the median in the NAI sample (reference). (B) Boxplot of SHAPE–MaP mutation frequencies for *B. subtilis* 16S and 23S rRNAs probed *in vivo*. Box plots span the interquartile range (from Q1 to Q3). The grey area spans from the median in the DMSO sample (control) to the median in the NAI sample (reference).

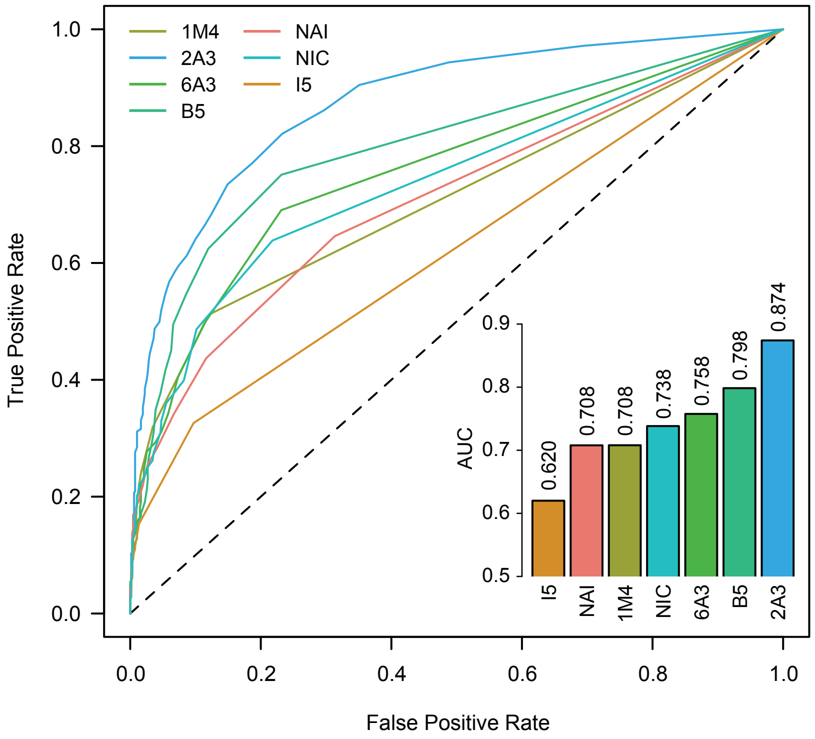


**Figure S4.** ROC curve for *in vivo* probing with all tested SHAPE reagents after subtracting DMSO mutation frequencies, calculated with respect to the accepted phylogenetically-inferred *E. coli* 16S and 23S rRNA structures from CRW. The inset reports the AUC for each compound.

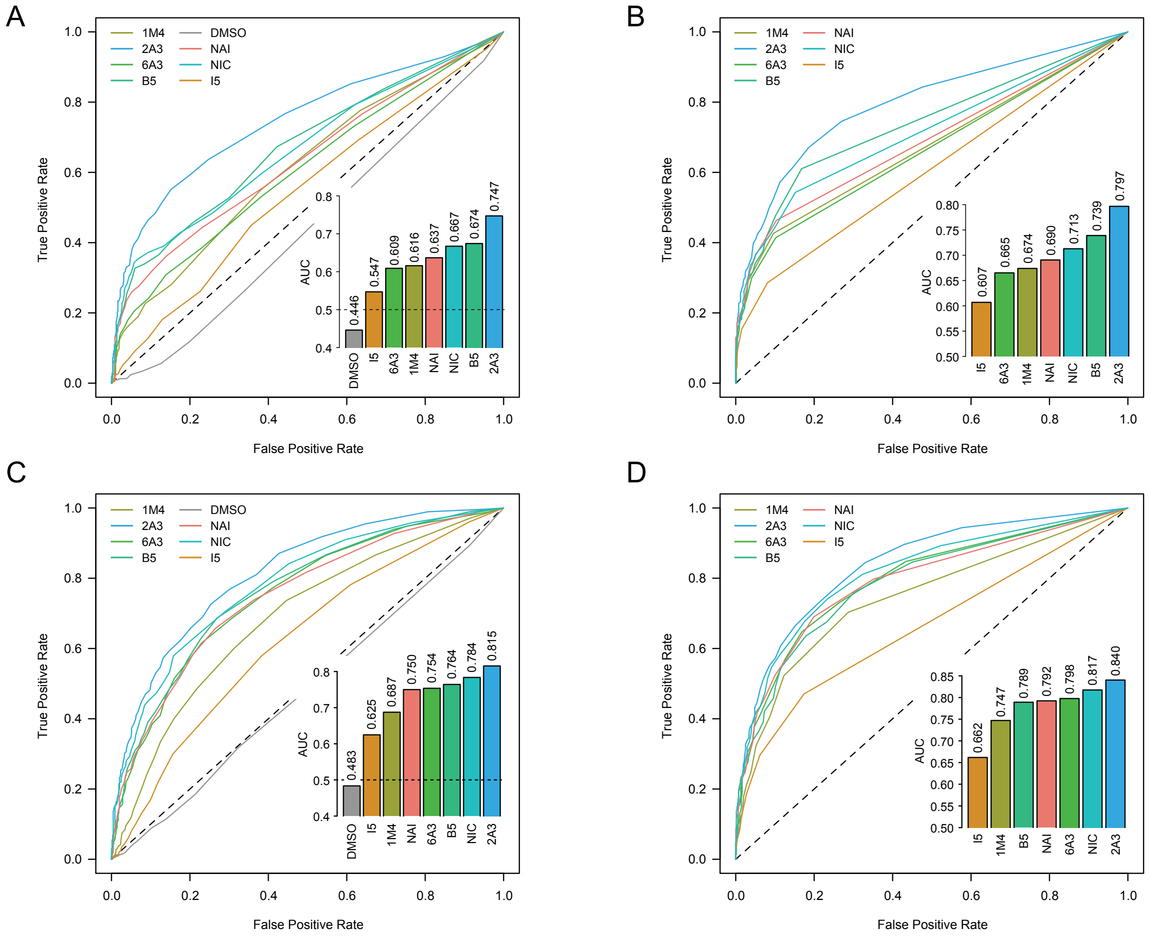


**Figure S5.** (A) and (B). ROC curve for *in vivo* probing with all tested SHAPE reagents before and after subtracting DMSO mutation frequencies, calculated on solvent-exposed residues in the crystal structure of the *B. subtilis* ribosome (PDB: 6HA1), with respect to the accepted phylogenetically-inferred *B. subtilis* 16S and 23S rRNA structures from CRW. (C) and (D). ROC curve for *in vivo* probing with all tested SHAPE reagents before and after subtracting DMSO mutation frequencies, calculated on solvent-exposed residues in the crystal structure of the *H. sapiens* ribosome (PDB: 4UG0), with respect to the accepted phylogenetically-inferred *H. sapiens* 18S and 28S rRNA structures from RNAcentral. The insets report the AUC for each compound.

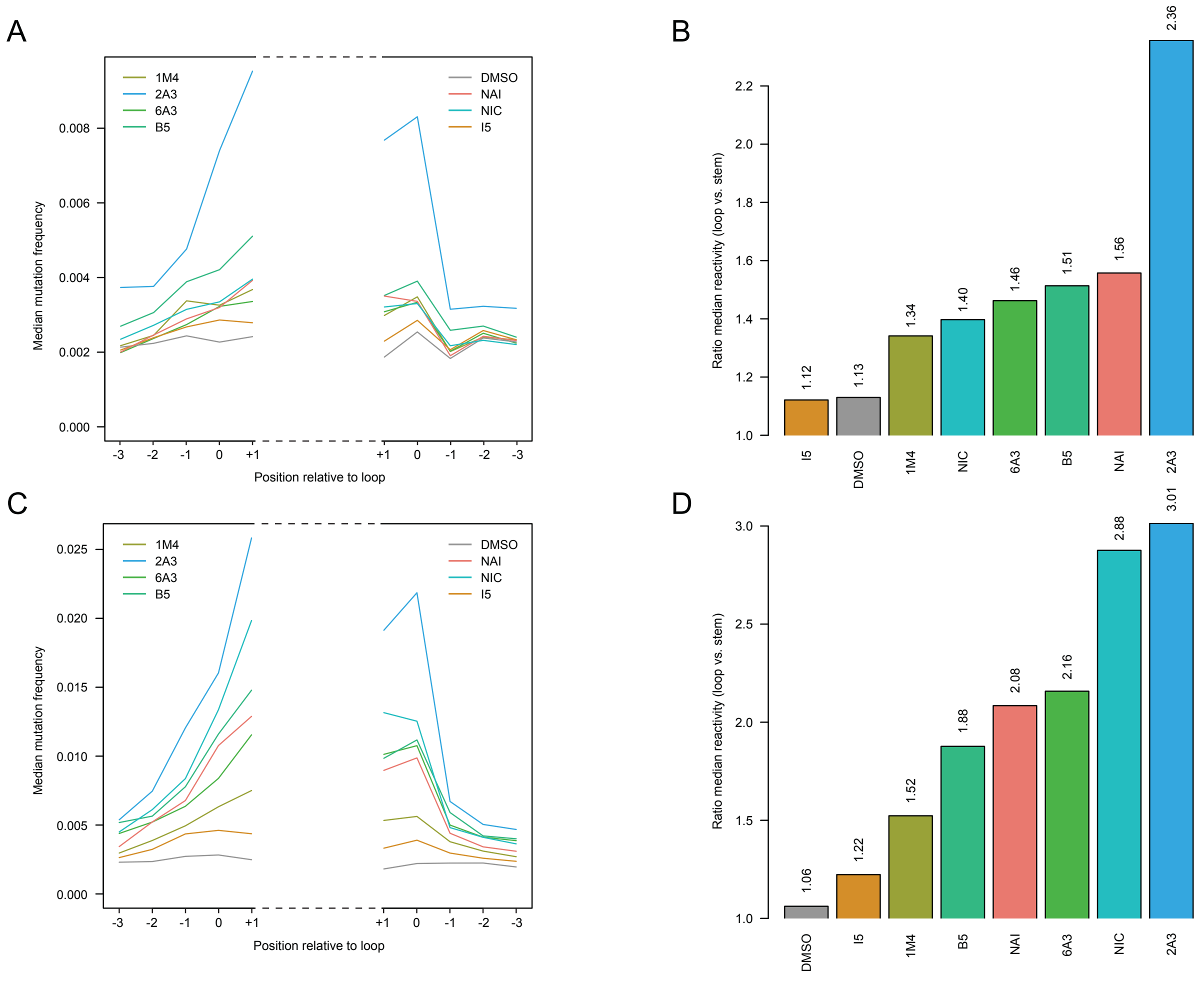


**Figure S6.** (A) Median *in vivo* SHAPE-MaP mutation frequencies across all stem-loops in the accepted *B. subtilis* 16S and 23S rRNA structures. Bases are numbered relatively to the loop. Positions -3 to -1 correspond to stem bases, while positions 0 and +1 correspond to loop bases. (B) Ratio between the median loop mutation frequency and the median stem mutation frequency, calculated on all stem-loops from A. (C) Median *in vivo* SHAPE-MaP mutation frequencies across all stem-loops in the accepted *H. sapiens* 18S and 28S rRNA structures. Bases are numbered relatively to the loop. Positions -3 to -1 correspond to stem bases, while positions 0 and +1 correspond to loop bases. (D) Ratio between the median loop mutation frequency and the median stem mutation frequency, calculated on all stem-loops from C.

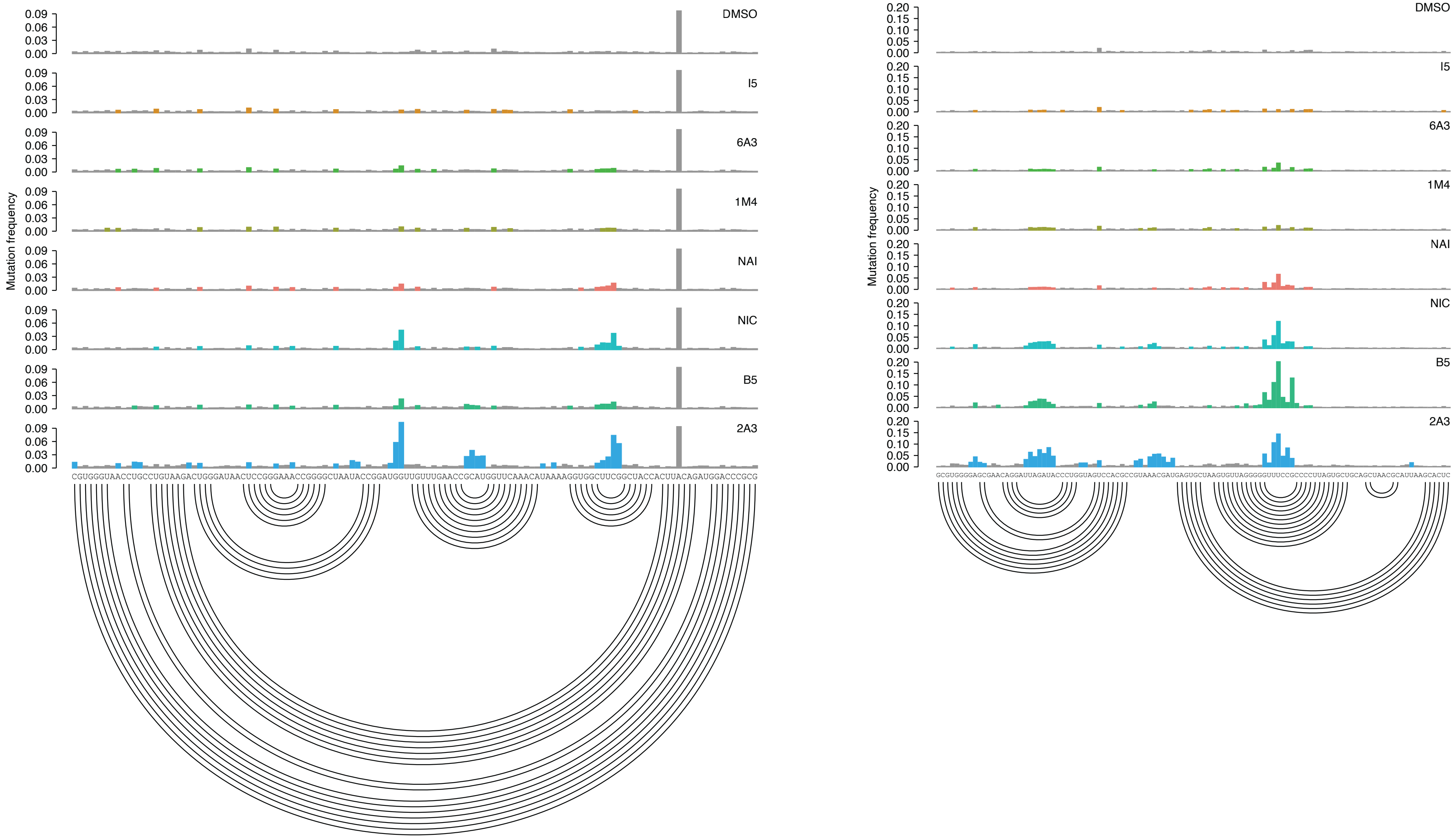


**Figure S7.** Sample of *in vivo* SHAPE−MaP mutation frequencies for all tested compounds across two regions, respectively spanning nucleotides 120 to 245 and nucleotides 777 to 888 of *B. subtilis* 16S rRNA. Colored bases are those whose mutation frequencies exceed by 2-folds the median mutation frequency in the analyzed region. The accepted structure is reported as an arc plot.

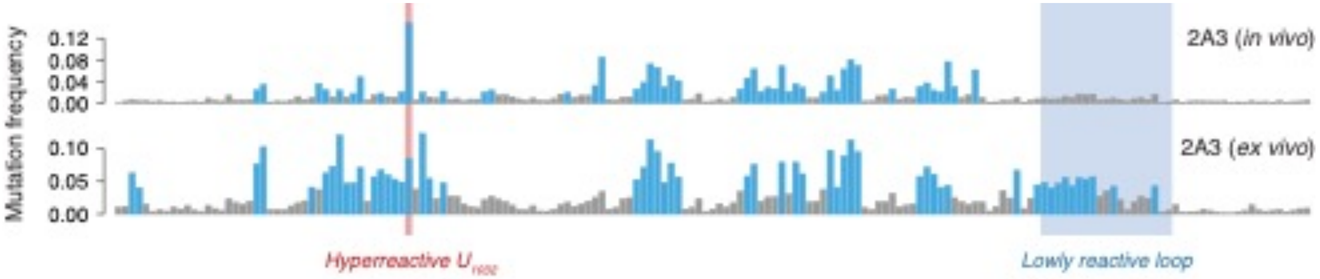


**Figure S8.** SHAPE–MaP mutation frequencies for 2A3 *ex vivo* and *in vivo* across a region spanning nucleotides 990 to 1162 of *E. coli* 23S rRNA (relative to Figure 3A). Colored bases are those whose mutation frequencies exceed by 2-folds the median mutation frequency in the analyzed region. The hyperreactive residue $U_{1032}$ and the unreactive stretch of 19 nucleotides are respectively marked in red and blue.

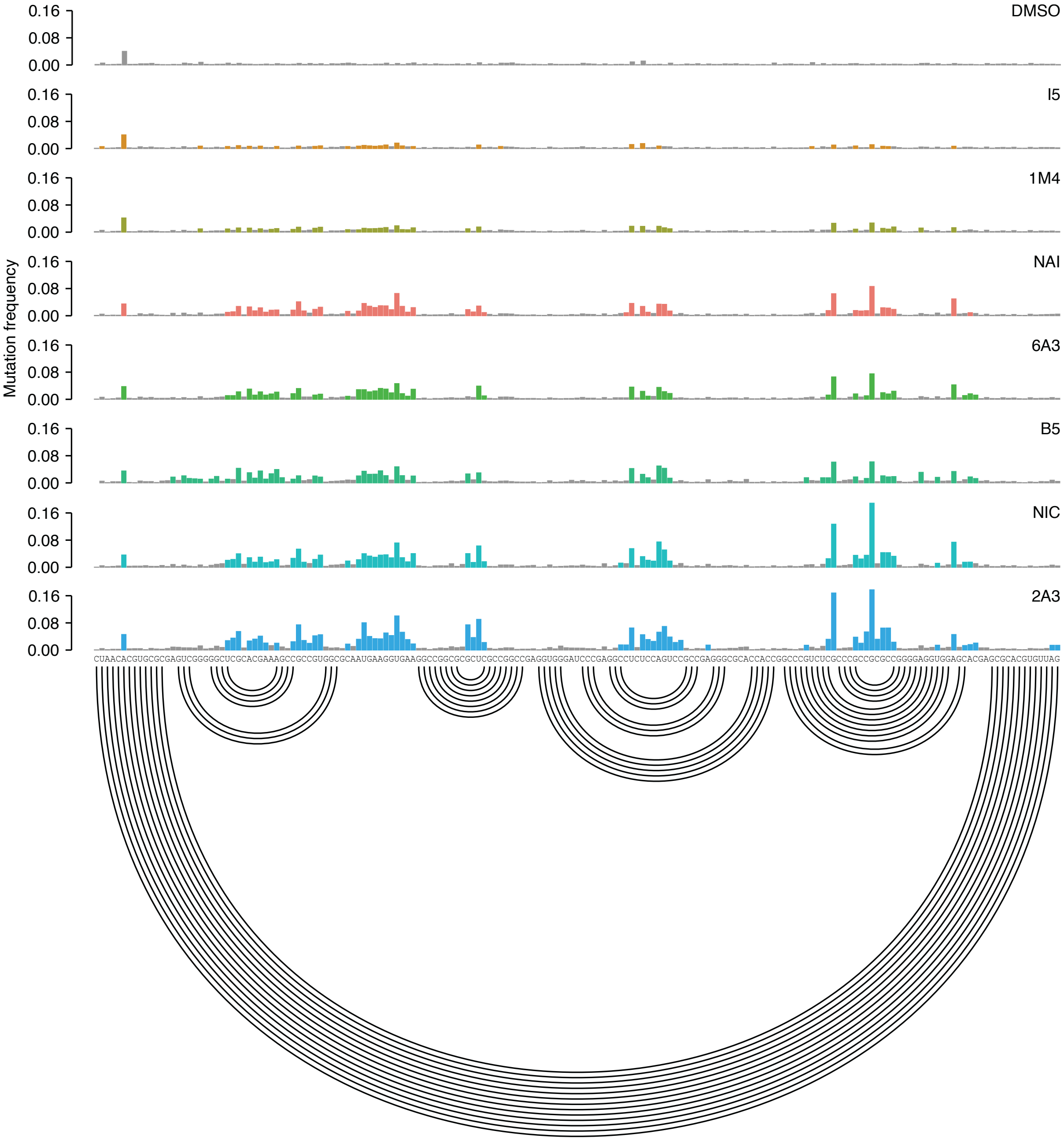


**Figure S9.** Sample of *in vivo* SHAPE−MaP mutation frequencies for all tested compounds across a region spanning nucleotides 1340 to 1516 of *H. sapiens* 28S rRNA. Colored bases are those whose mutation frequencies exceed by 2-folds the median mutation frequency in the analyzed region. The accepted structure is reported as an arc plot.

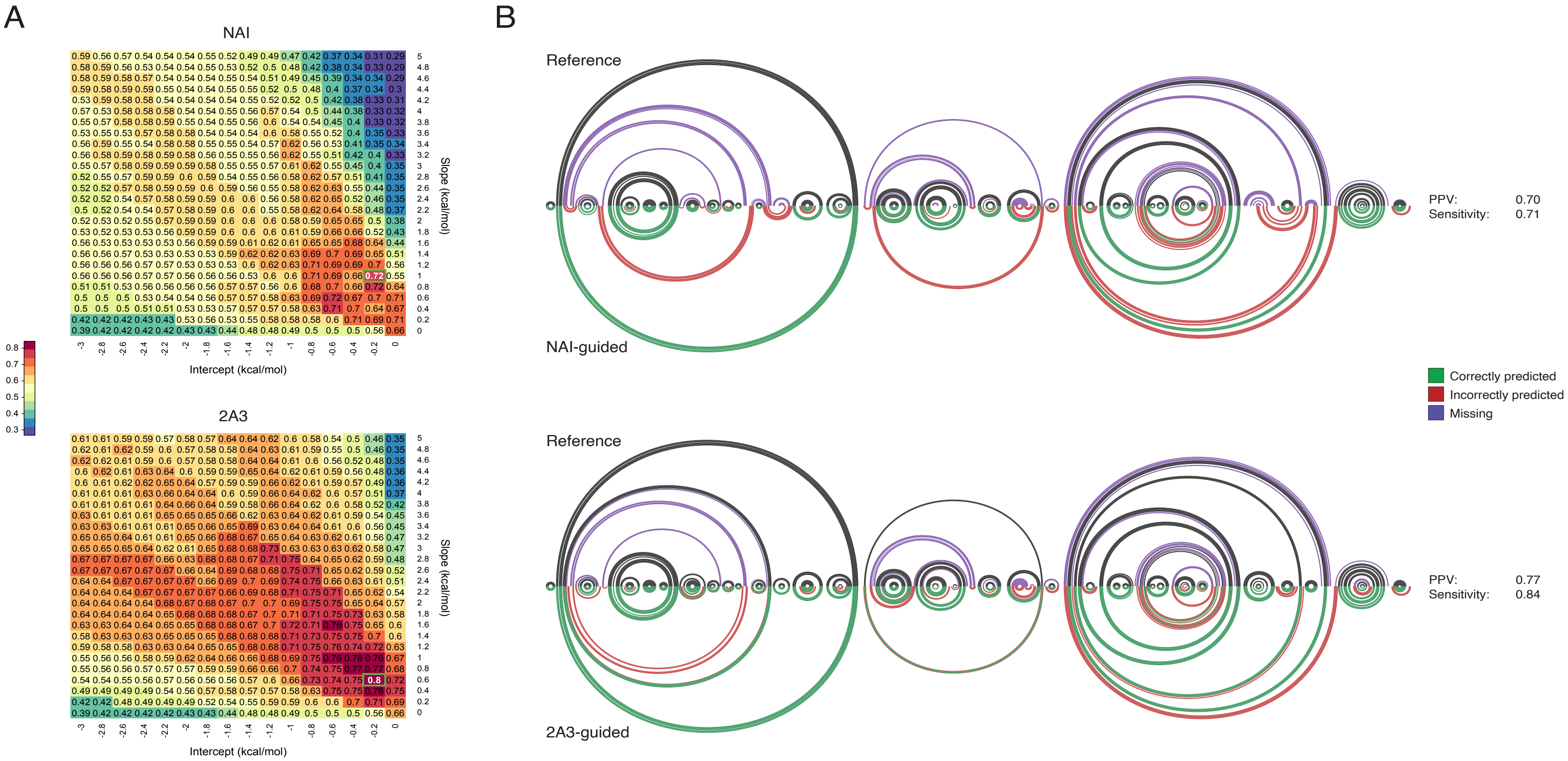


**Figure S10.** (A) Grid search (jackknifing) of optimal slope/intercept value pairs for *B. subtilis* 16S and 23S rRNAs *in vivo* probing data for NAI and 2A3, performed using ViennaRNA 2.0. Values represent the geometric mean of sensitivity and PPV for the secondary structures predicted using each slope/intercept value pair. The chosen value pair is boxed in green. (B) Arc plot comparison of *B. subtilis* 16S rRNA reference structure (top), and structure inferred using either NAI-derived or 2A3-derived restraints (bottom). Black/green arcs correspond to correctly predicted base-pairs, violet arcs to non-predicted base-pairs and red arcs to mispredicted base-pairs. PPV and sensitivity for each prediction are indicated.

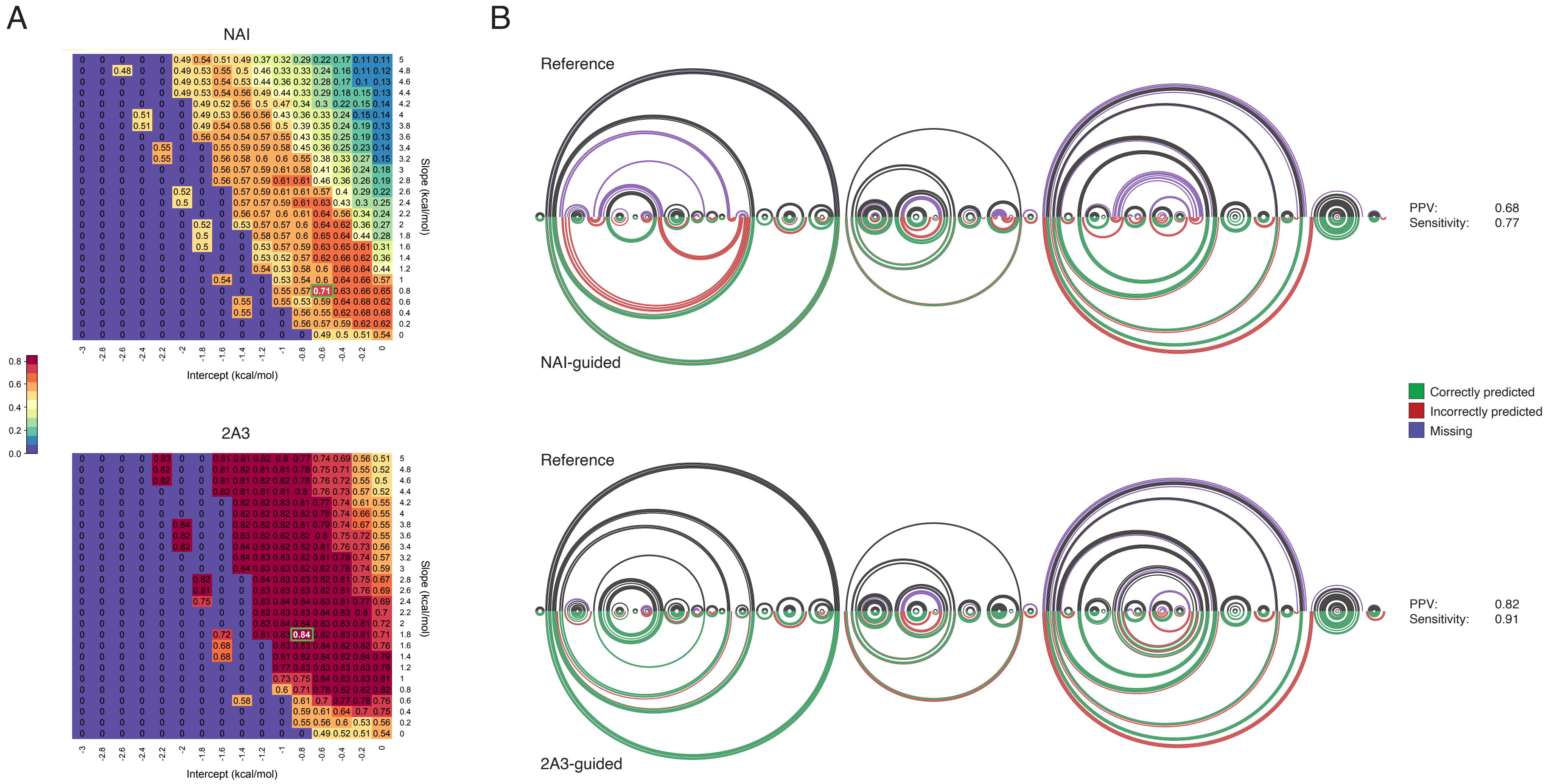


**Figure S11.** (A) Grid search (jackknifing) of optimal slope/intercept value pairs for *E. coli* 16S and 23S rRNAs *in vivo* probing data for NAI and 2A3, performed using RNAstructure. Values represent the geometric mean of sensitivity and PPV for the secondary structures predicted using each slope/intercept value pair. Pairs with geometric mean equal to 0, produced an empty structure (no base-pairs). The chosen value pair is boxed in green. (B) Arc plot comparison of *E. coli* 16S rRNA reference structure (top), and structure inferred using either NAI-derived or 2A3-derived restraints (bottom). Black/green arcs correspond to correctly predicted base-pairs, violet arcs to non-predicted base-pairs and red arcs to mispredicted base-pairs. PPV and sensitivity for each prediction are indicated.

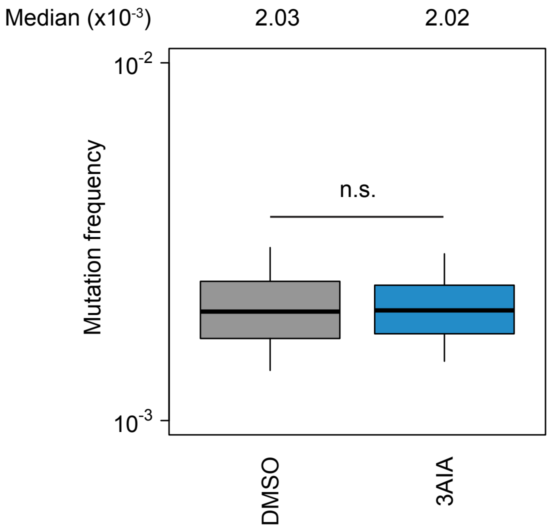


**Figure S12.** Boxplot of SHAPE–MaP mutation frequencies for *E. coli* 16S and 23S rRNAs probed *in vivo*. Box plots span the interquartile range (from Q1 to Q3). 3AIA is unable to probe RNA *in vivo* as shown by absence of any significant difference with respect to the DMSO control ($P = 0.2$, Wilcoxon rank sum test).

| Electrophile | 25 °C | 37 °C |
|---|---|---|
| NIC | 30 minutes | 8 minutes |
| 2A3 | 260 minutes | 60 minutes |
| 6A3 | 510 minutes | 300 minutes |
| B5 | 70 minutes | 30 minutes |
| 1M4 | 900 minutes | 290 minutes |
| 3AIA | 44 minutes | 14 minutes |
| NAI | 300 minutes | 120 minutes |

**Table S1.** Half-life data for SHAPE electrophiles in pH 7.4 PBS 1X at 25°C and 37°C

#### Chemical characterization and half-life determination

**Nicotinic acid imidazolide (NIC)**

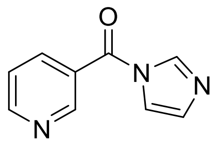


NIC for SHAPE experiment was obtained following the general procedure described in the Materials and Methods section. For the analytical sample, a portion of the crude mixture was diluted into dichloromethane and extracted 3X with saturated sodium bicarbonate solution. The organic layer was dried over $MgSO_4$ and concentrated under reduced pressure.

$C_9H_7N_3O$, $^1H$ NMR (300 MHz, DMSO-d6): δ 7.20 (s, 1H,), 7.65 (dd, 1H), 7.73 (s, 1H), 8.22 (dt, 1H), 8.25 (s, 1H), 8.90 (d, 1H), 8.98 (s, 1H) ppm, $^{13}C$ NMR (300 MHz, DMSO-d6): δ 118.24, 123.74, 128.17, 130.67, 137.46, 138.51, 149.98, 153.46, 164.77 ppm, HRMS: Compound not identified

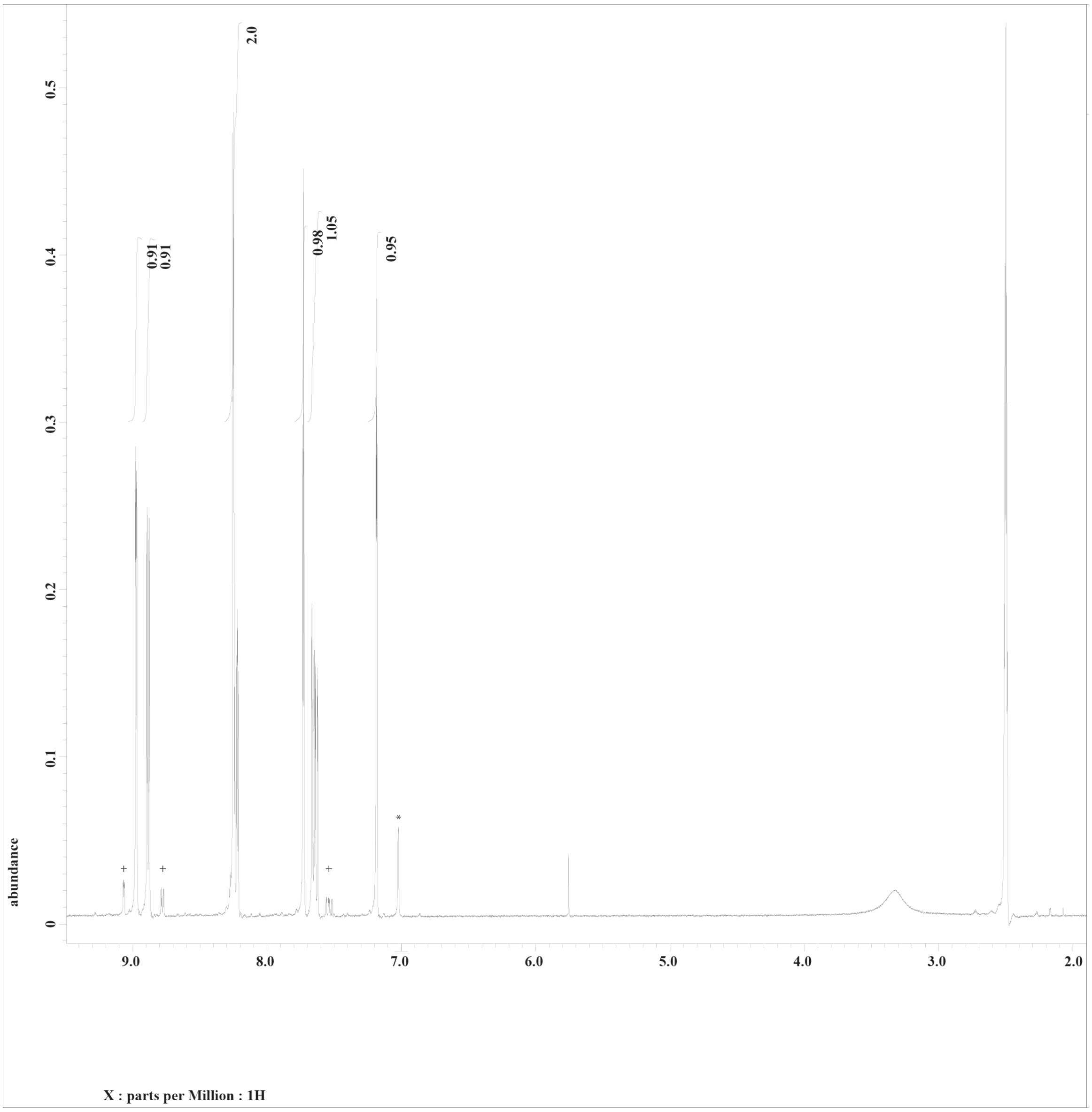


[1]H NMR of NIC in DMSO-d6. + signifies hydrolyzed product protons and * shows imidazole protons.

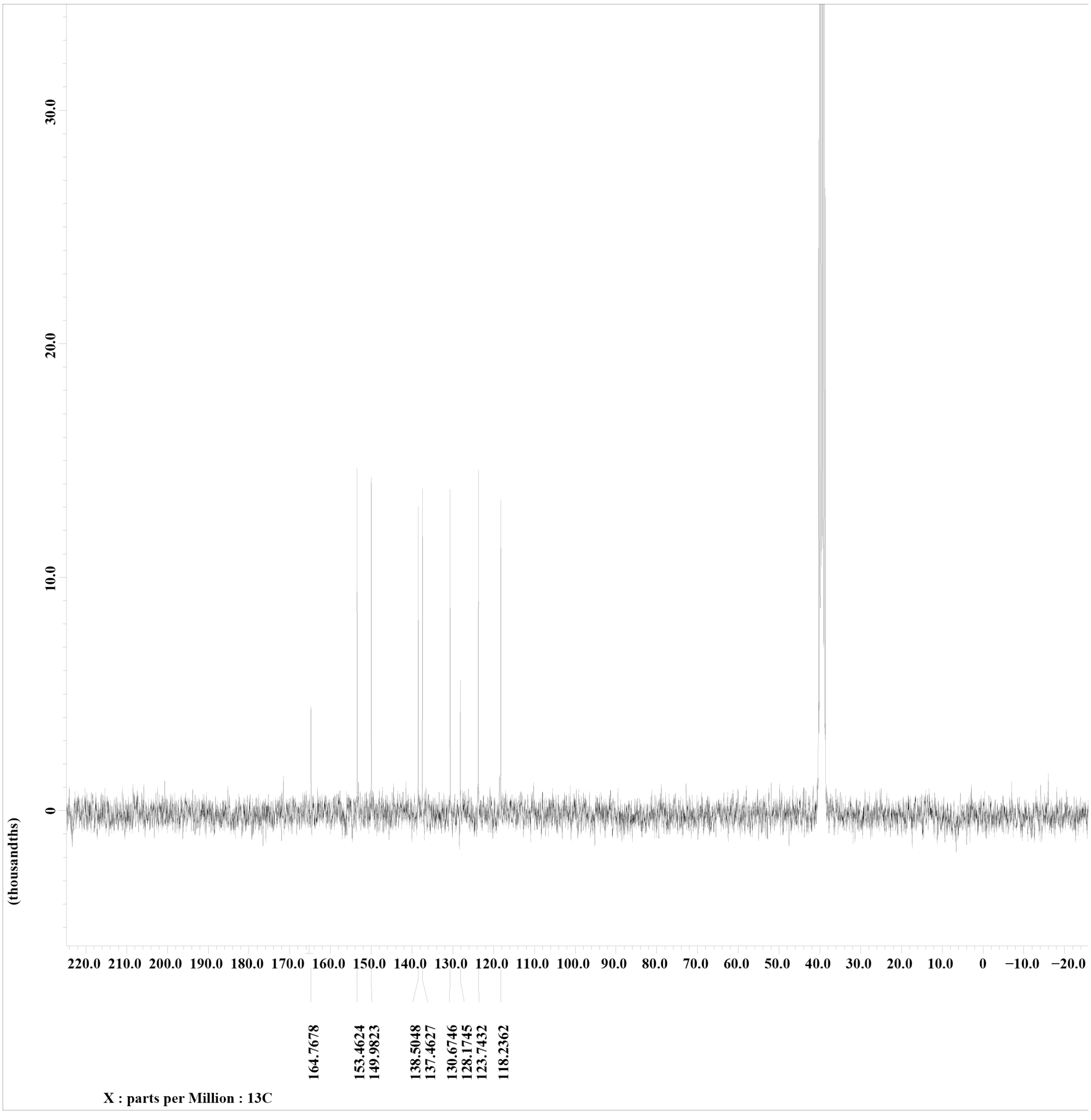


$^{13}$C-NMR of NIC in DMSO-d6.

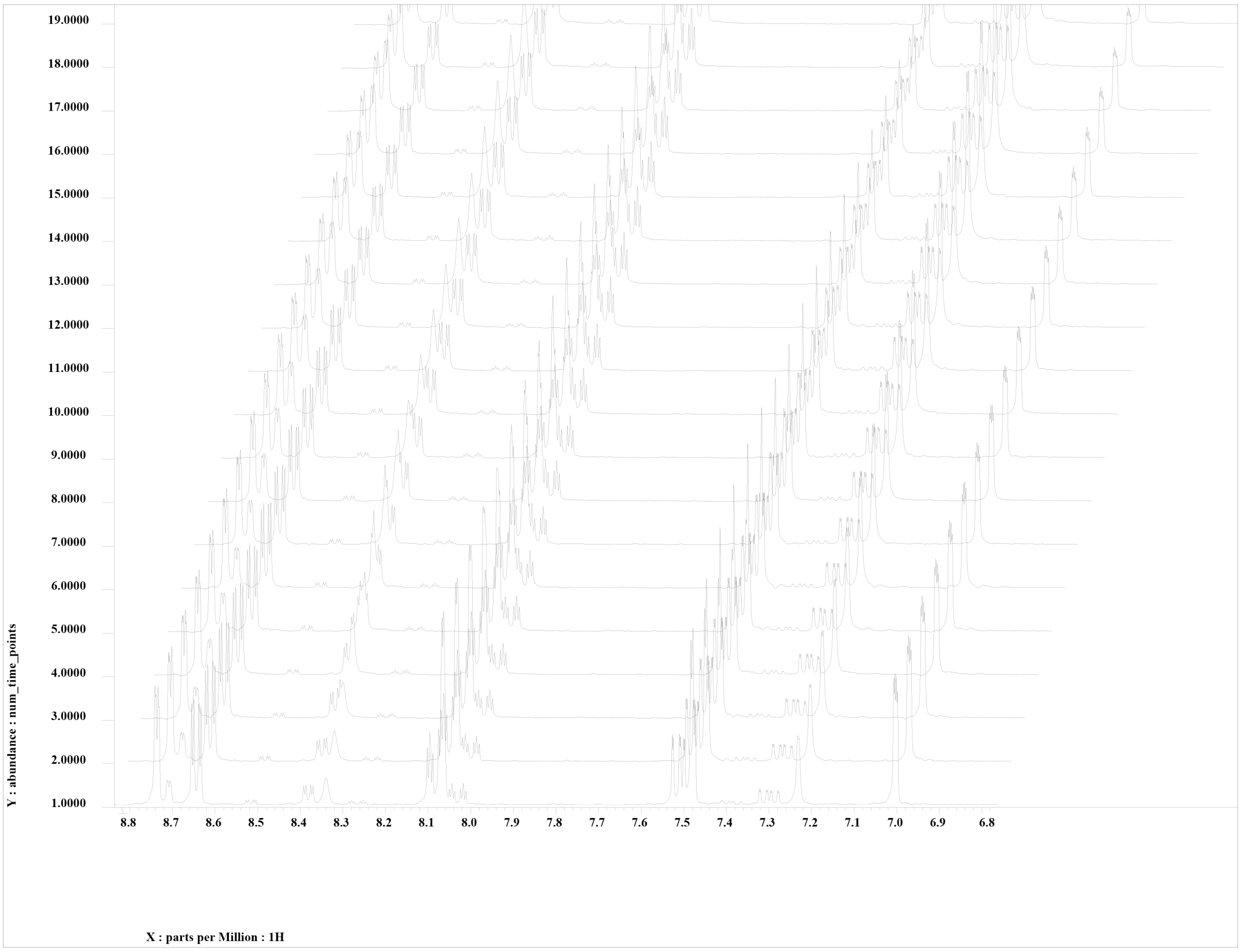


Hydrolysis experiment of NIC at 25°C. Scan 1 occurred at 5 minutes, each subsequent scan was performed in 2 minutes intervals. Monitoring the disappearance of product peaks at 8.81 and 7.48 ppm.

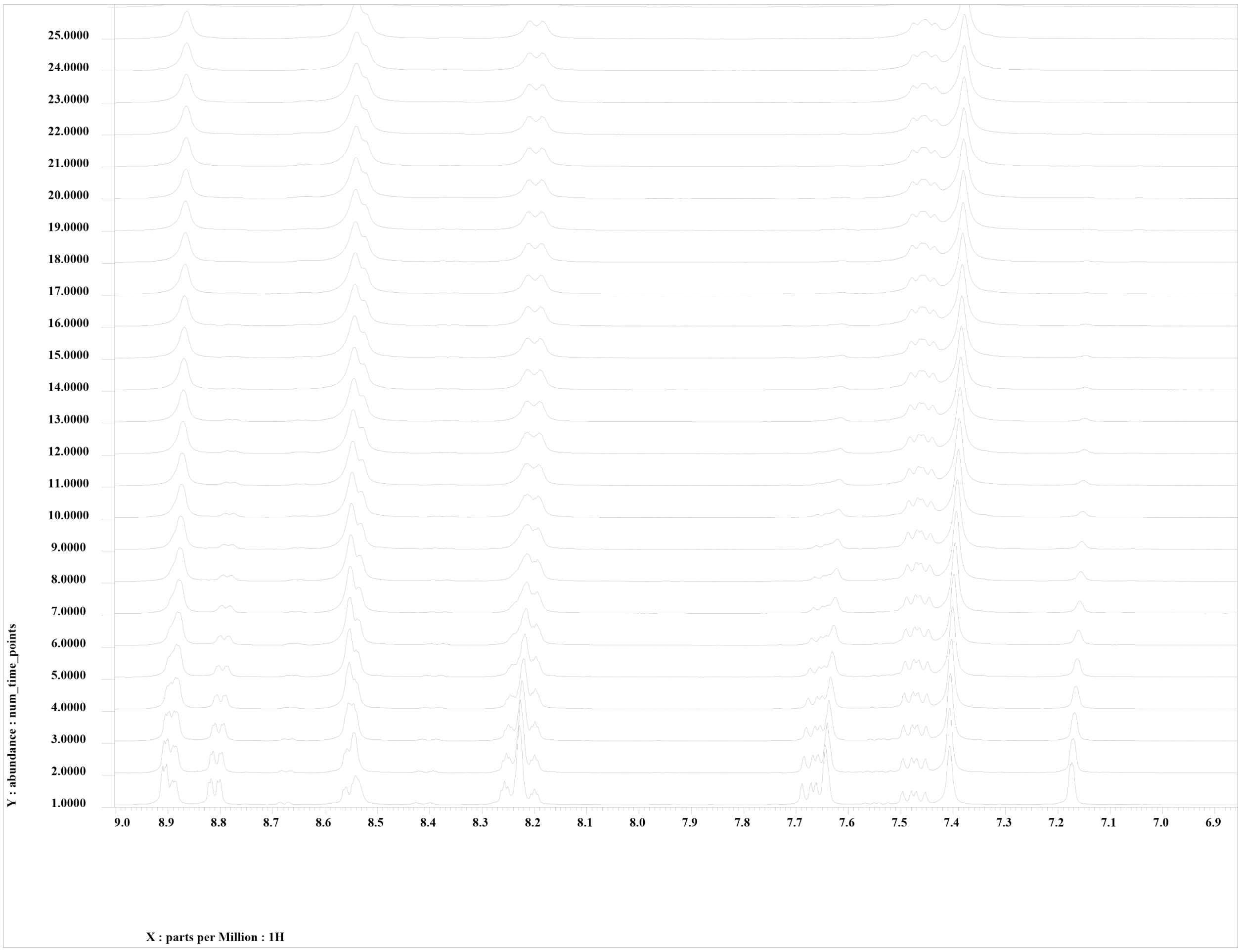


Hydrolysis experiment of NIC at 37°C. Scan 1 occurred at 5 minutes, each subsequent scan was performed in 2 minutes intervals. Monitoring the disappearance of product peaks at 8.81 and 7.48 ppm.

.

**2-aminopyridine-3-carboxylic acid imidazolide (2A3)**

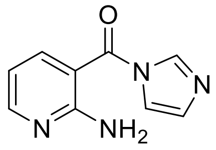

2A3 was obtained following the general procedure described in the Materials and Methods section, but using MeCN in place of DMSO. For the analytical sample, a portion of the crude mixture was diluted into dichloromethane and extracted 3X with saturated sodium bicarbonate solution. The organic layer was dried over $MgSO_4$ and concentrated under reduced pressure.

$C_9H_8N_4O$, $^1H$ NMR (300 MHz, DMSO-d6): δ 6.69 (t, 1H,), 7.04(s, 2H), 7.15 (s, 1H), 7.65 (s, 1H), 7.76 (d, 1H), 8.18 (s, 1H), 8.28 (dd, 1H) ppm, $^{13}C$ NMR (300 MHz, DMSO-d6): δ 107.11, 111.69, 118.57, 130.03, 138.09, 140.52, 154.15, 159.20, 165.78 ppm, HRMS Found: 189.0771 (M+H) m/z.

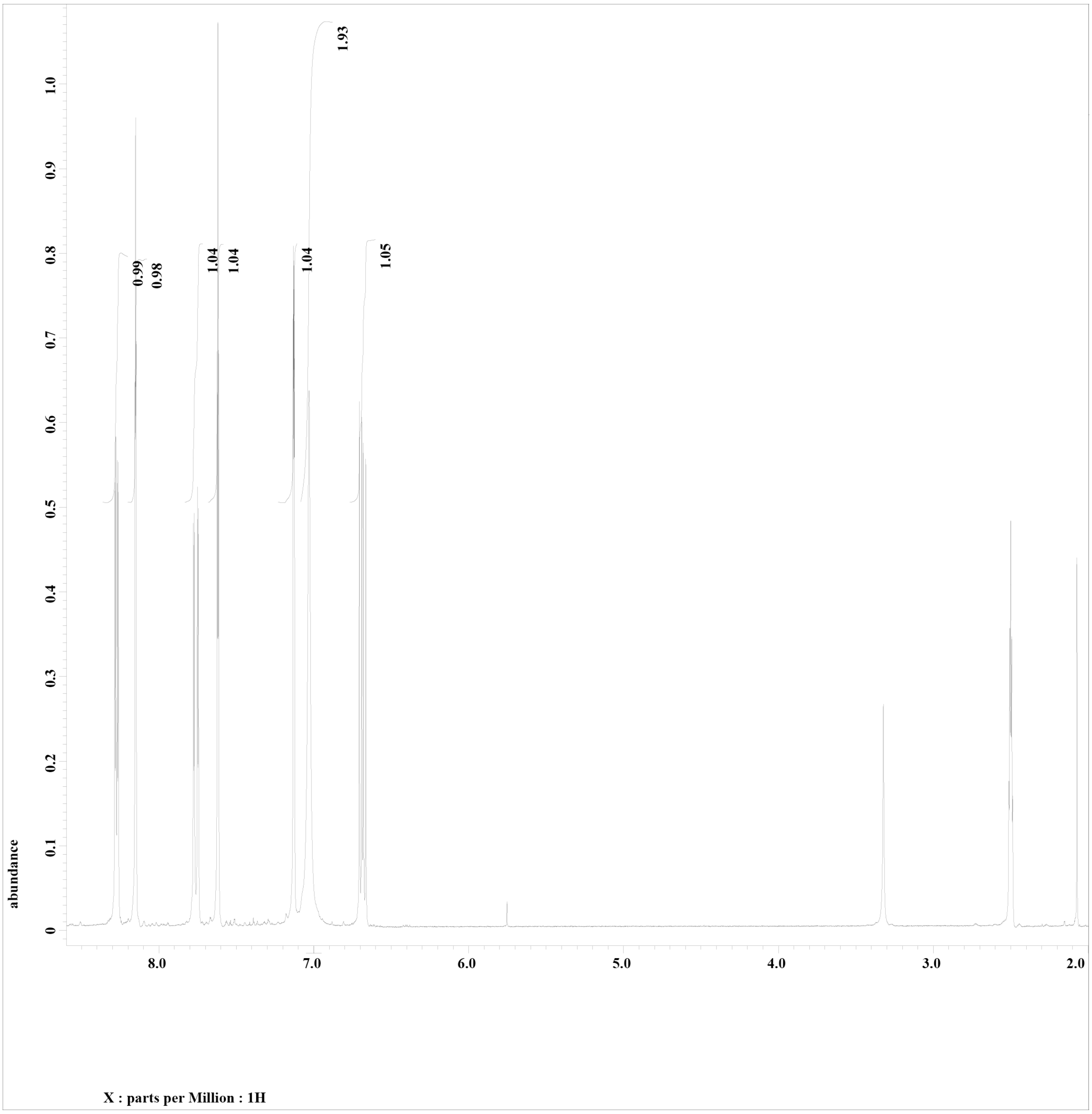


[1]H NMR of 2A3 in DMSO-d6.

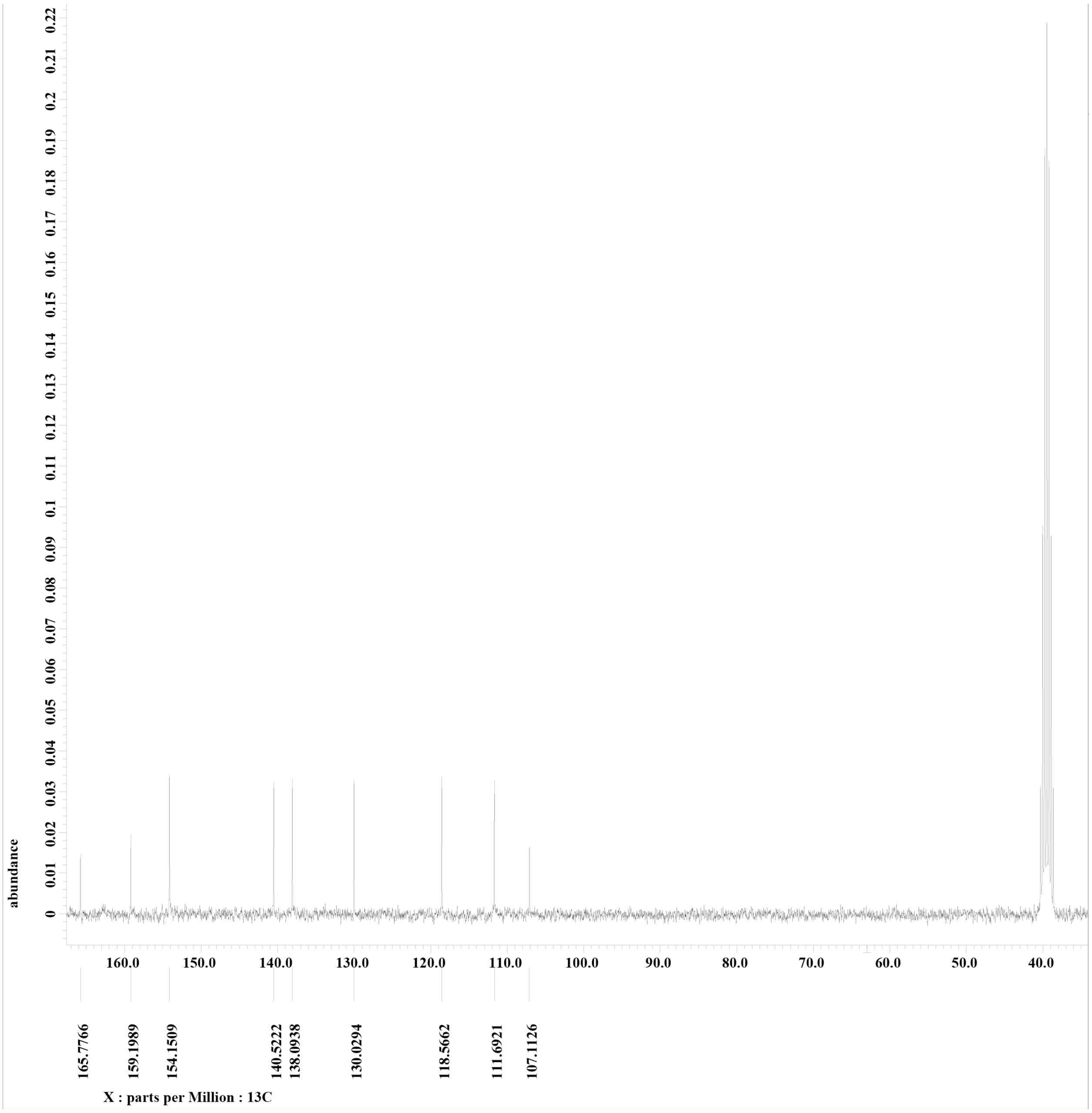


$^{13}$C-NMR of 2A3 in DMSO-d6.

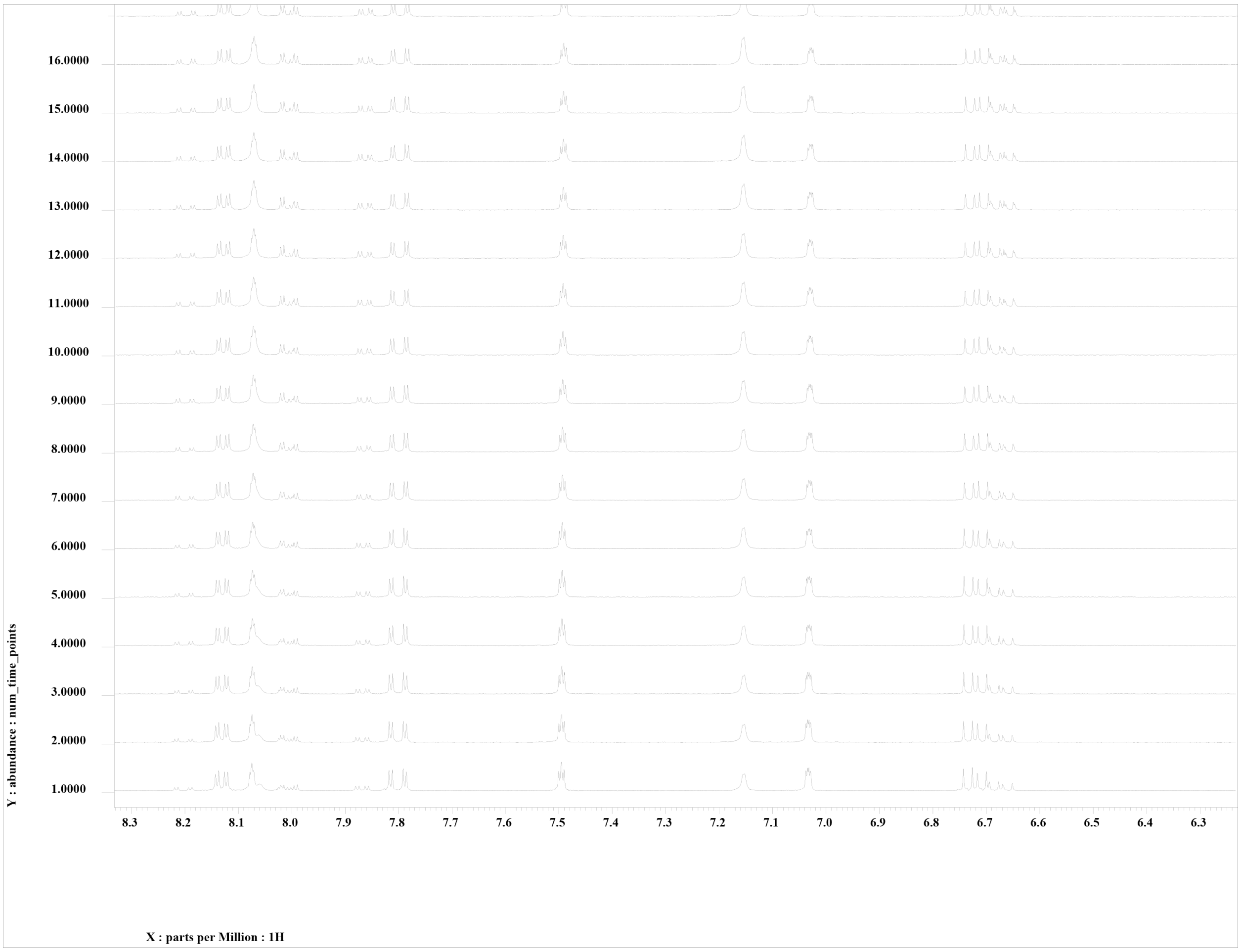


Hydrolysis experiment of 2A3 at 25°C. Showing 85 minutes to 185 minutes, where each scan was performed in 5 minutes intervals. Monitoring the disappearance of product peaks at 8.81 and 7.48 ppm.

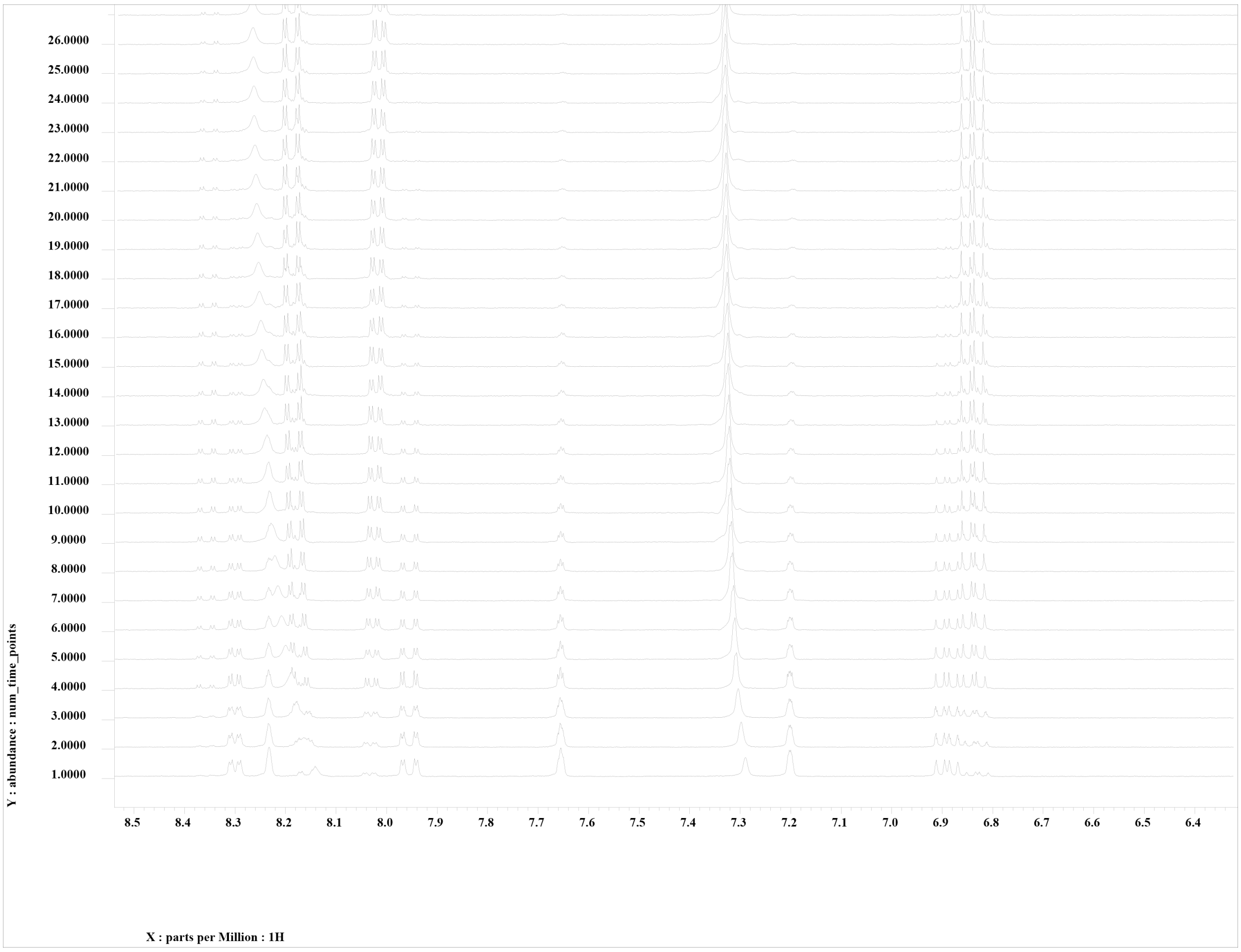


Hydrolysis experiment of 2A3 at 37°C. Showing 85 minutes to 185 minutes, where each scan was performed in 5 minutes intervals. Monitoring the disappearance of product peaks at 8.81 and 7.48 ppm.

**6-aminopyridine-3-carboxylic acid imidazolide (6A3)**

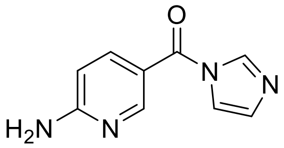

6A3 was obtained following the general procedure described in the Materials and Methods section. For the analytical sample, a portion of the crude mixture was diluted into dichloromethane and extracted 3X with saturated sodium bicarbonate solution. The organic layer was dried over $MgSO_4$ and concentrated under reduced pressure.

$C_9H_8N_4O$, $^1H$ NMR (300 MHz, DMSO-d6): δ 6.55 (d, 1H,), 7.12 (s, 1H), 7.16 (s, 2H), 7.78 (s, 1H), 7.82 (dd, 1H), 8.24 (s, 1H), 8.42 (d, 1H) ppm, $^{13}C$ NMR (300 MHz, DMSO-d6): δ 107.43, 114.11, 118.55, 129.90, 137.95, 138.73, 152.63, 162.92, 164.21 ppm, HRMS Found: 189.0771 (M+H) m/z.

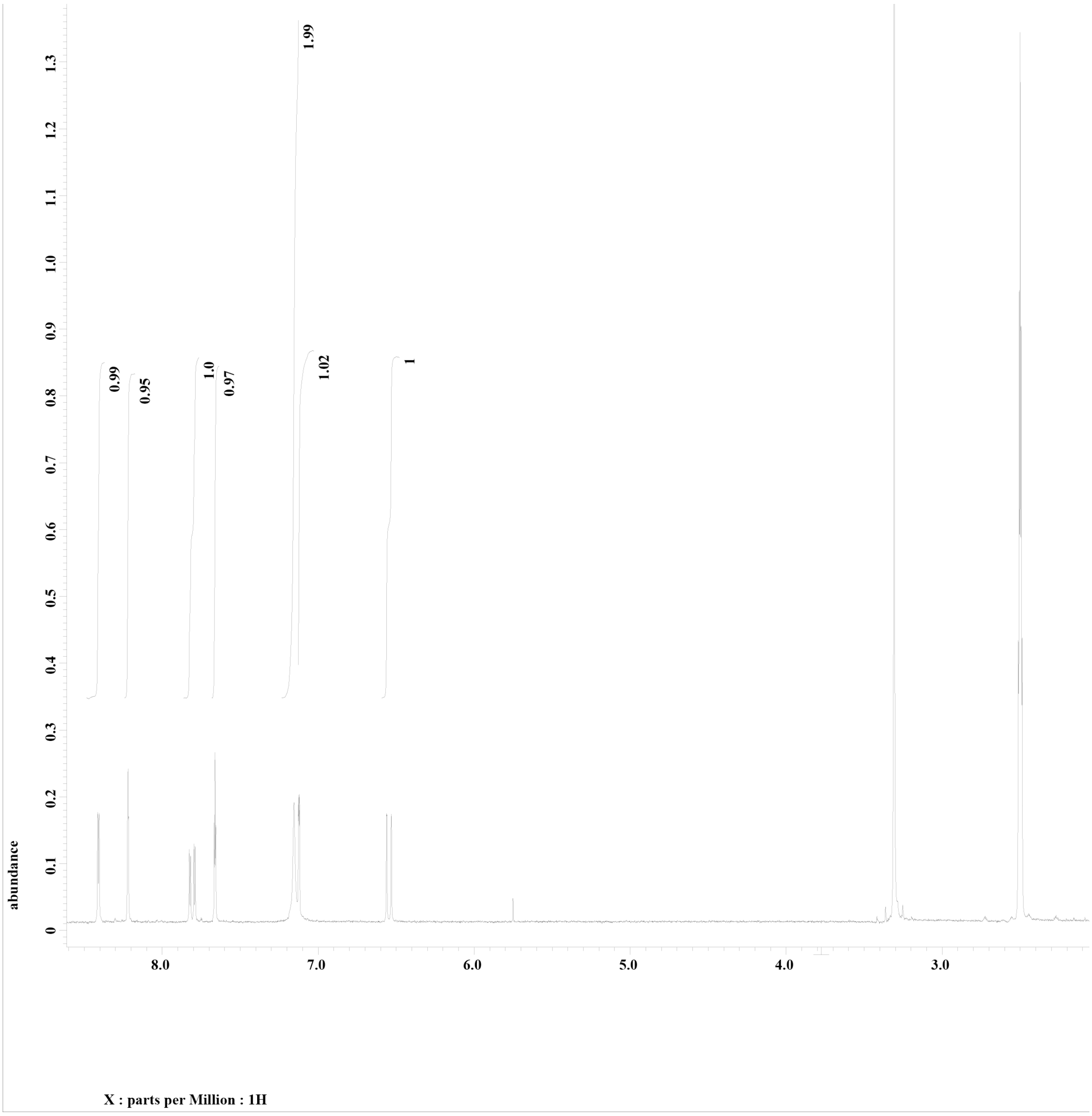


$^1$H NMR of 6A3 in DMSO-d6

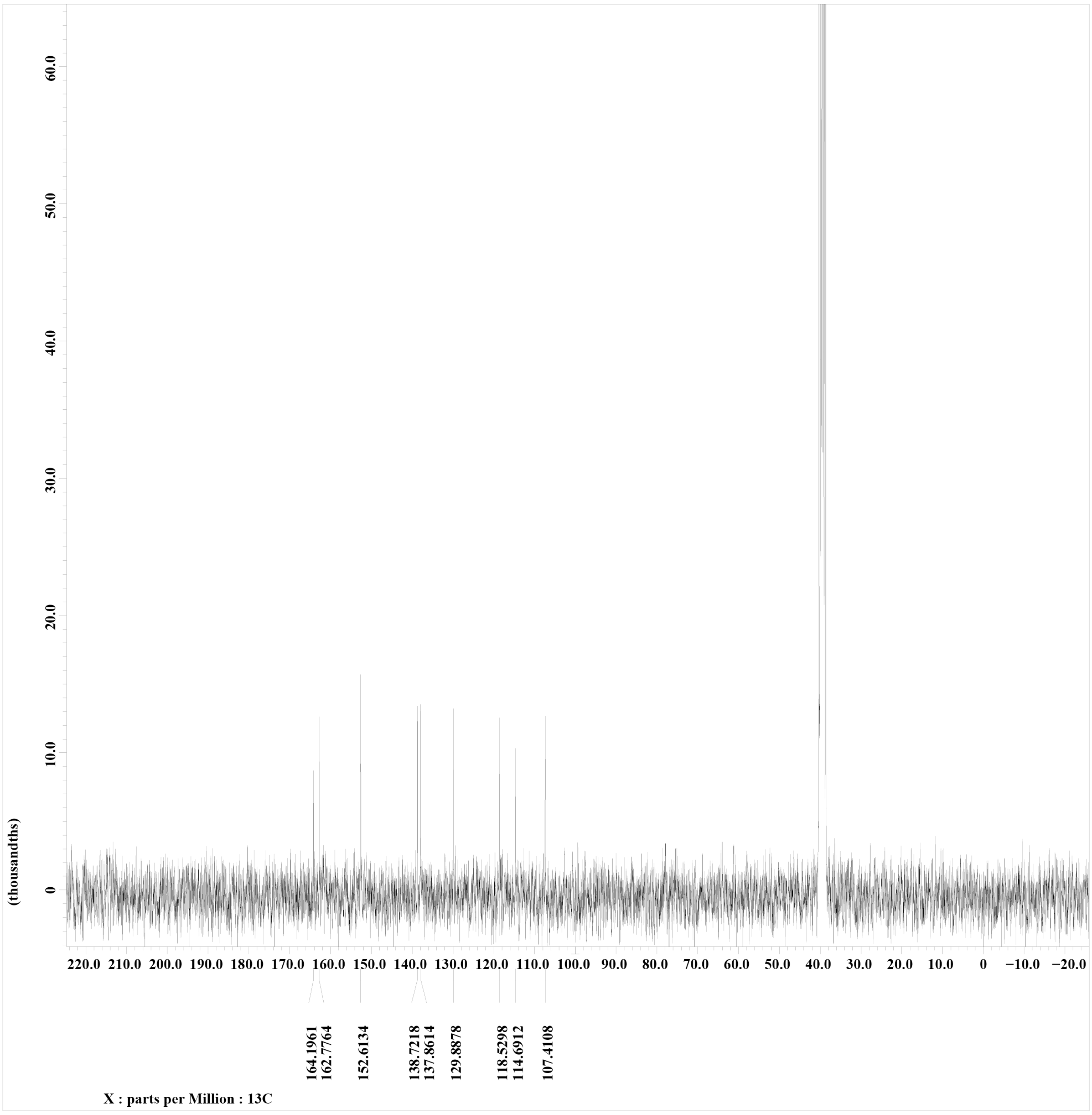


$^{13}C$ NMR of 6A3 in DMSO-d6

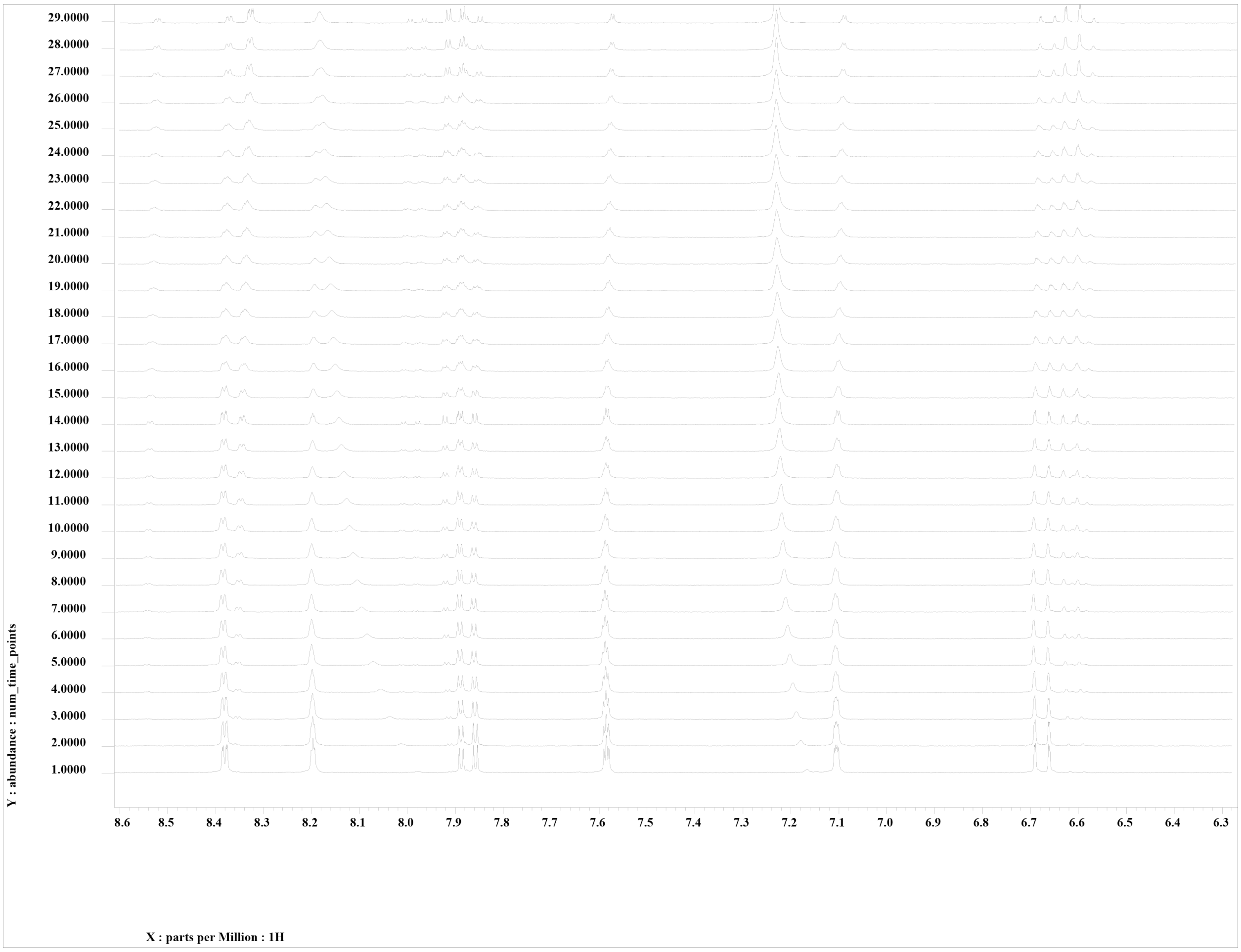


Hydrolysis experiment of 6A3 at 25°C. Showing 5 minutes to 905 minutes, where each scan was performed in 30 minutes intervals. Monitoring the disappearance of product peaks at 8.39 ppm and appearance at 8.36 ppm.

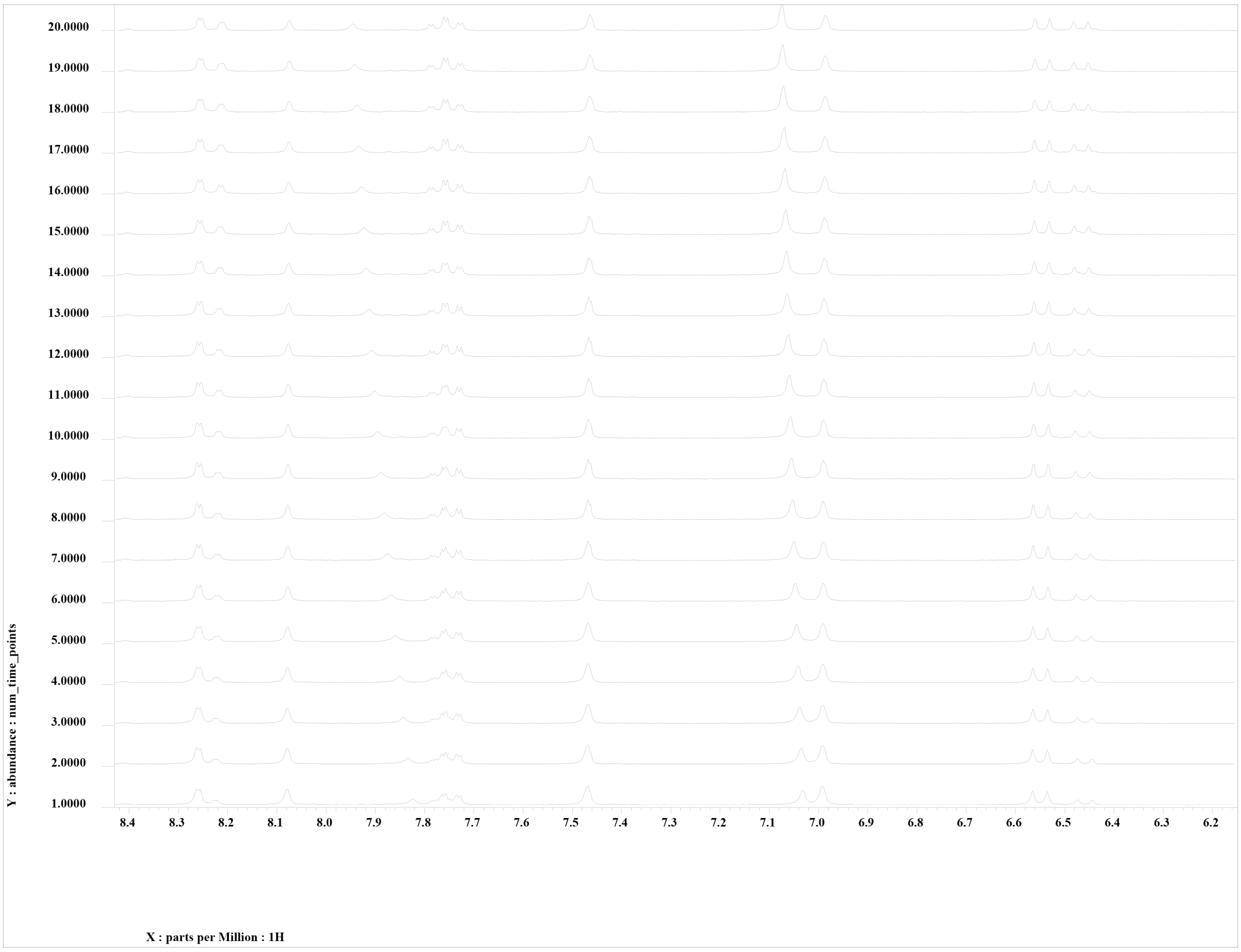


Hydrolysis experiment of 6A3 at 37°C. Showing 5 minutes to 905 minutes, where each scan was performed in 30 minutes intervals. Monitoring the disappearance of product peaks at 8.39 ppm and appearance at 8.36 ppm.

**Benzotriazole-5-carboxylic acid imidazolide (B5)**

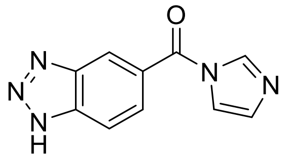

B5 was obtained following the general procedure described in the Materials and Methods section. Analytical samples were taken directly from the reaction mixture.

$C_{10}H_7N_5O$, $^1H$ NMR (300 MHz, DMSO-d6): δ 7.18 (s, 1H,), 7.72 (s, 1H), 7.84 (d, 2H), 8.05 (d, 1H), 8.26 (s, 1H), 8.45 (s, 1H) ppm, $^{13}C$ NMR (300 MHz, DMSO-d6): δ 114.61, 118.64, 119.80, 126.70, 128.14, 130.49, 138.72, 139.05, 139.90, 165.99 ppm, HRMS: Calc. Compound not identified

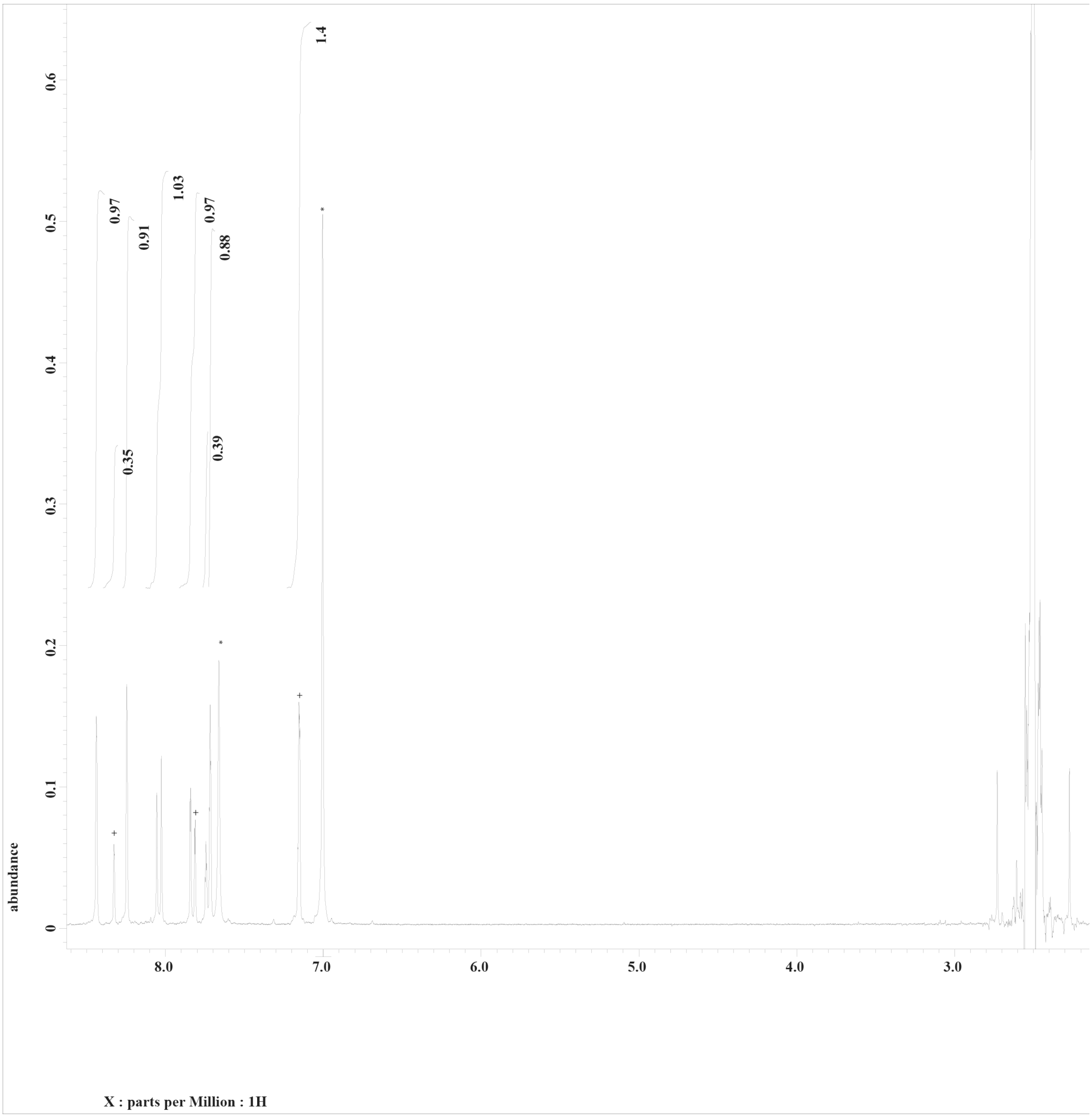


[1]H NMR of B5 in DMSO. Integration of three protons from excess CDI shown with +, where one falls under product peak leading to the high integration. Two signals from imidazole shown with *.

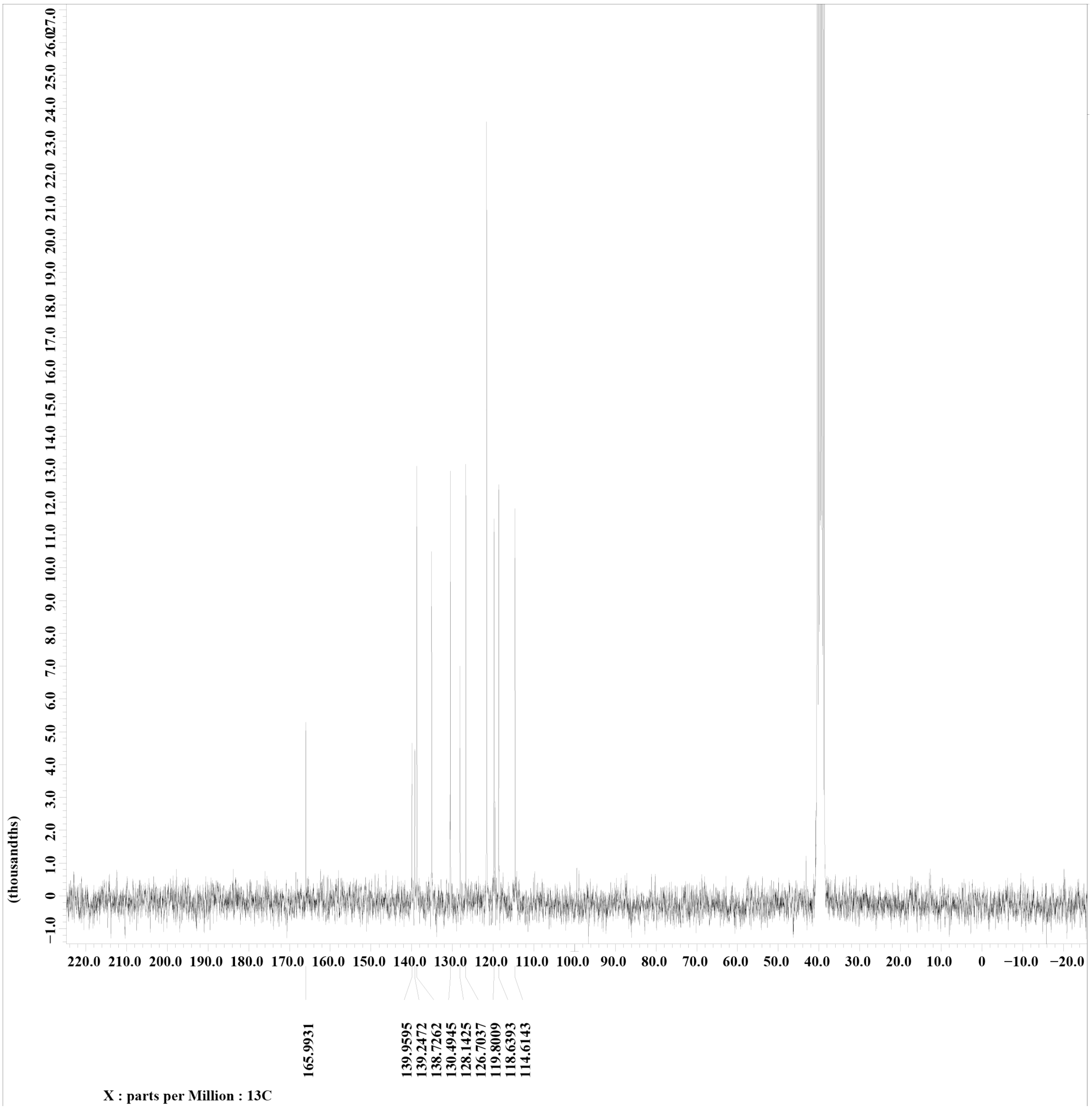


$^{13}$C-NMR of B5 in DMSO-d6. Liberated imidazole carbons shown with *.

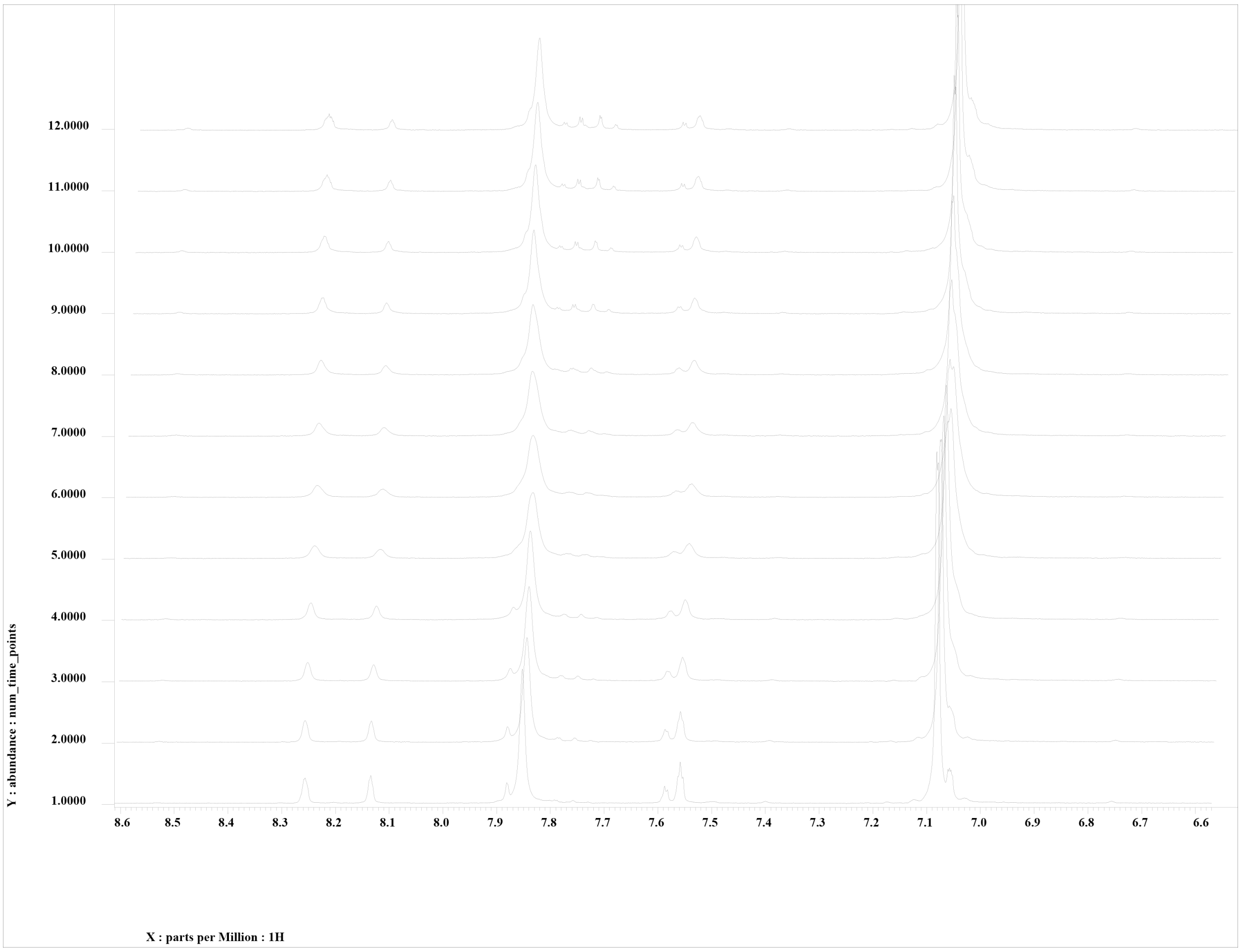


Hydrolysis experiment of B5 at 25°C. Showing from 5 minutes to 125 minutes, where each scan was performed in 10 minutes intervals. Monitoring the disappearance of product peaks at 8.28 ppm.

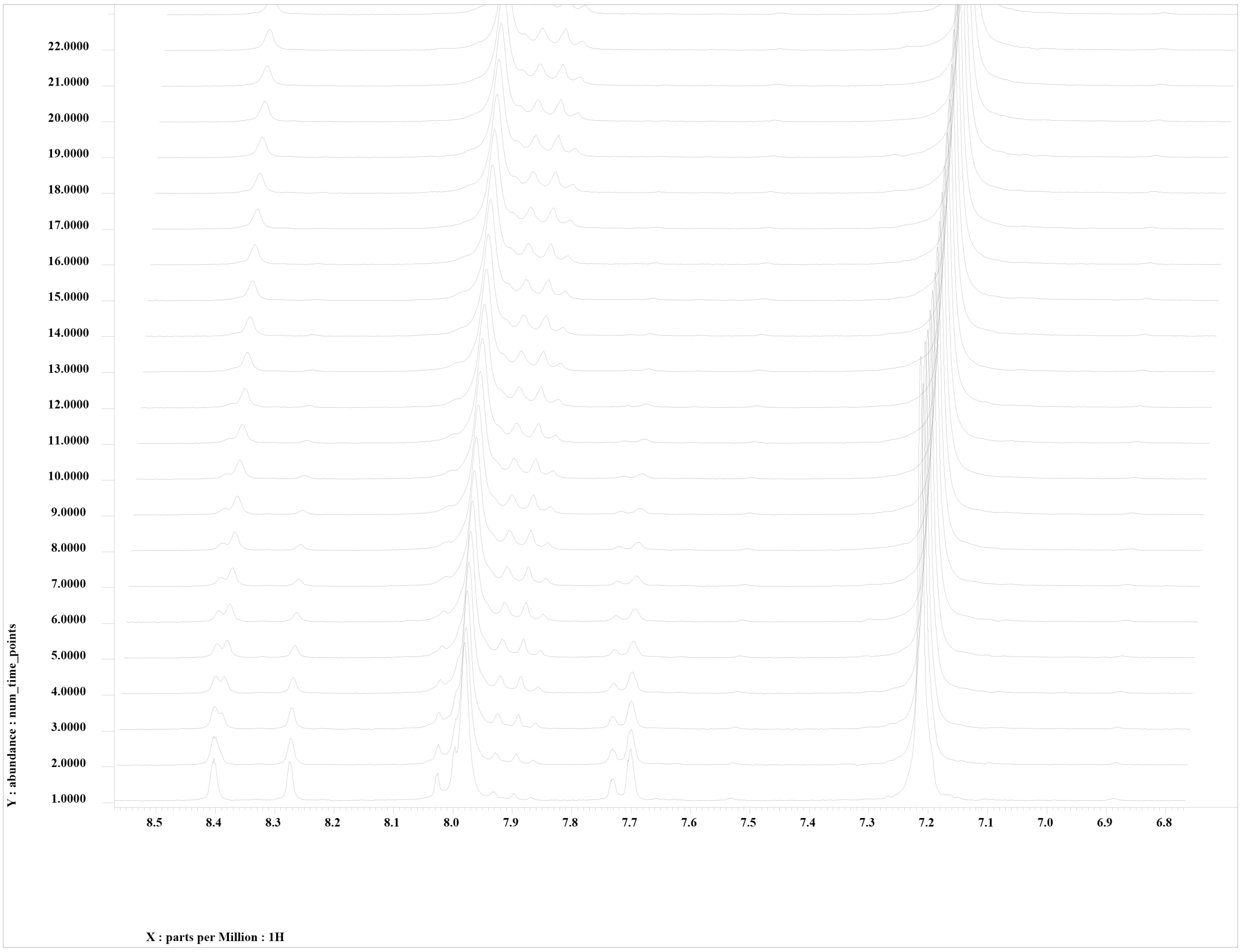


Hydrolysis experiment of B5 at 37°C. Showing from 5 minutes to 125 minutes, where each scan was performed in 10 minutes intervals. Monitoring the disappearance of product peaks at 8.28 ppm.

**1-methylimidazole-4-carboxylic acid imidazolide (1M4)**

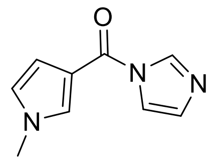

1M4 was obtained following the general procedure described in the Materials and Methods section. For the analytical sample, a portion of the crude mixture was diluted into dichloromethane and extracted 3X with saturated sodium bicarbonate solution. The organic layer was dried over $MgSO_4$ and concentrated under reduced pressure.

$C_9H_9N_3O$, $^{1}H$ NMR (300 MHz, DMSO-d6): δ 3.78 (s, 3H,), 7.10 (s, 1H), 7.94 (s, 1H), 8.08 (s, 1H), 8.32 (s, 1H), 9.00 (s, 1H) ppm, $^{13}C$ NMR (300 MHz, DMSO-d6): δ 33.65, 117.82, 129.53, 131.51, 133.22, 138.51, 138.22, 138.51, 139.93, 158.31 ppm, HRMS Found: 177.0771 (M+H) m/z.

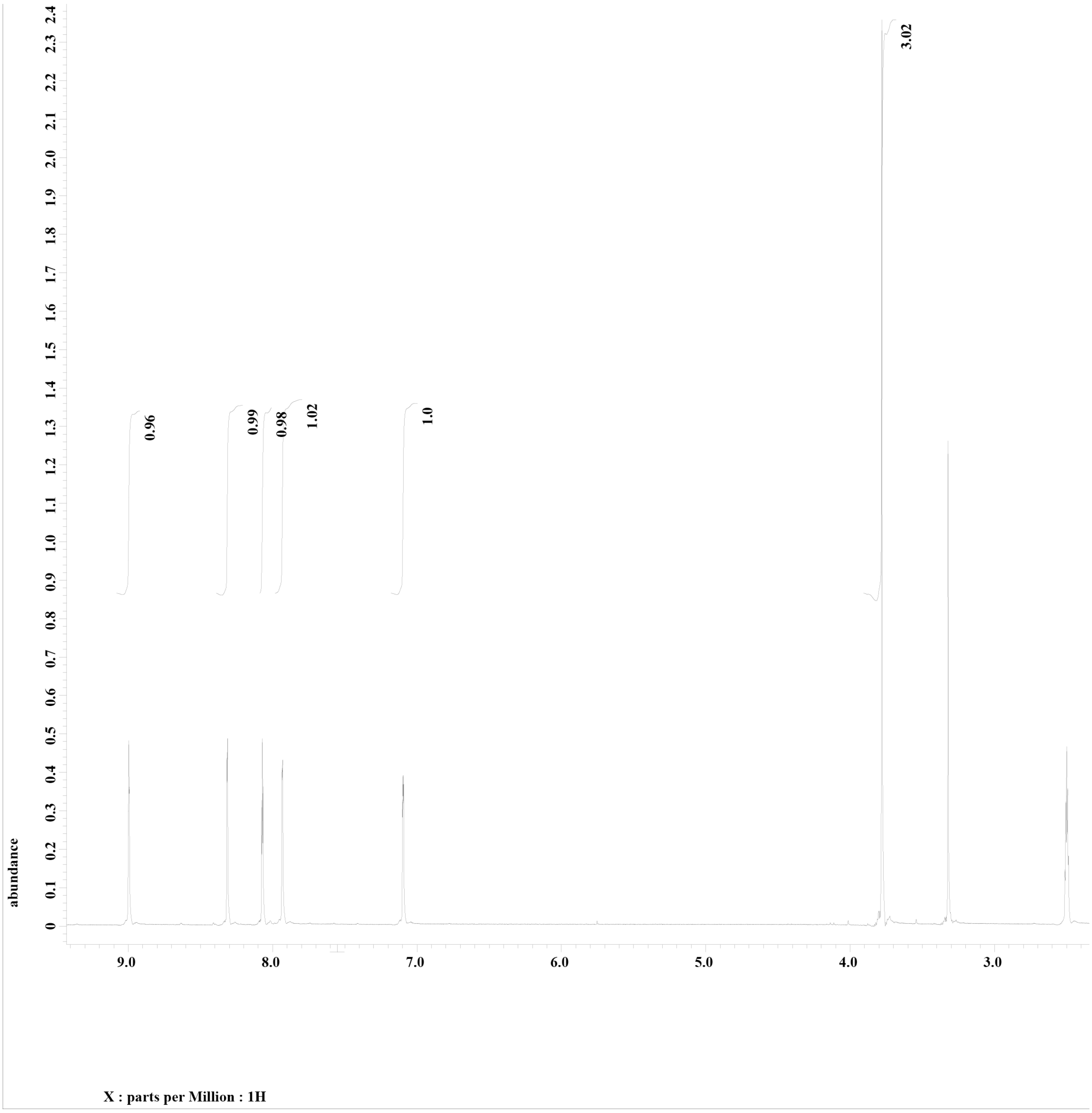


$^1$H NMR of 1M4in DMSO-d6

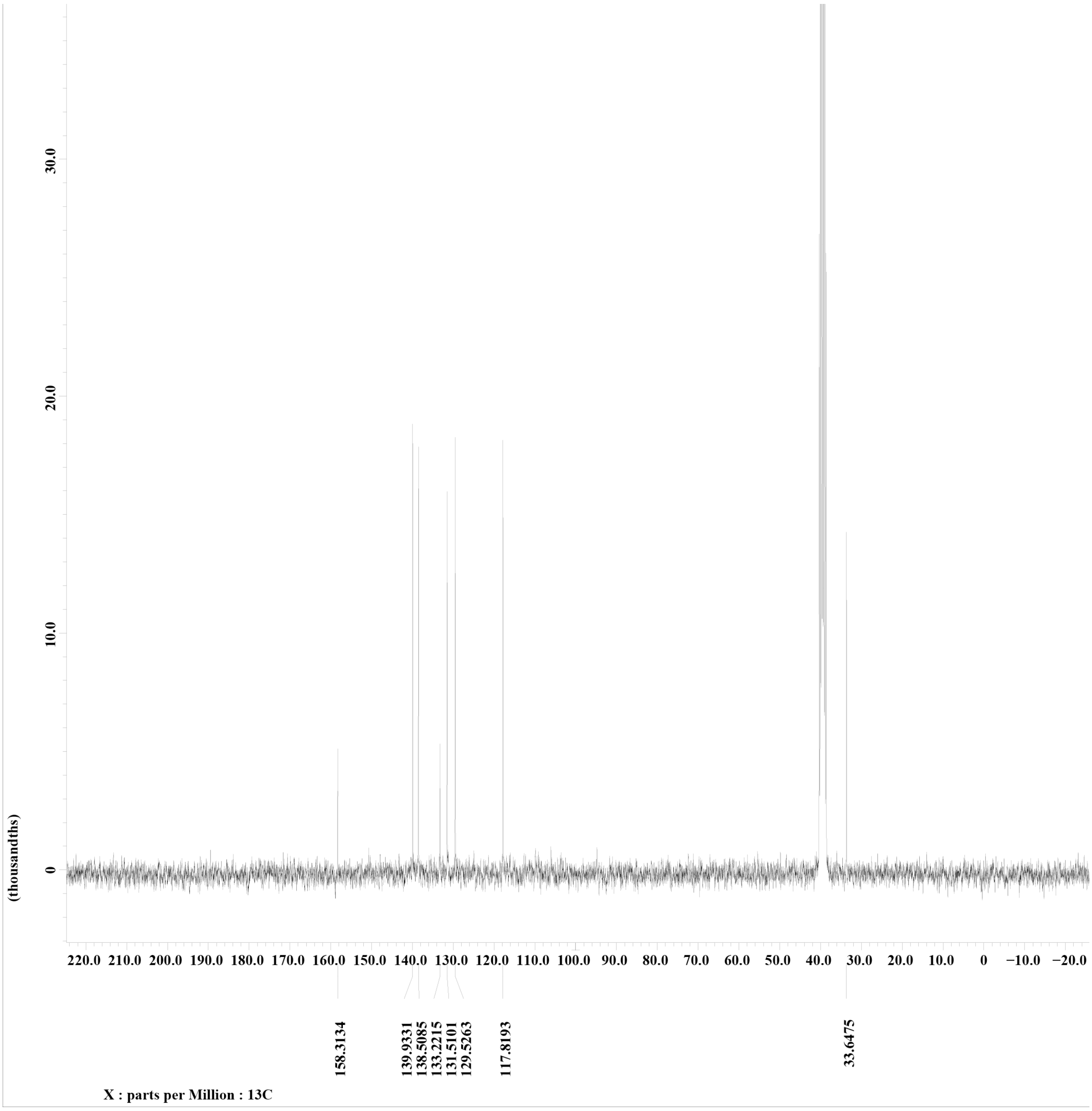


$^{13}C$ NMR of 1M4 in DMSO-d6.

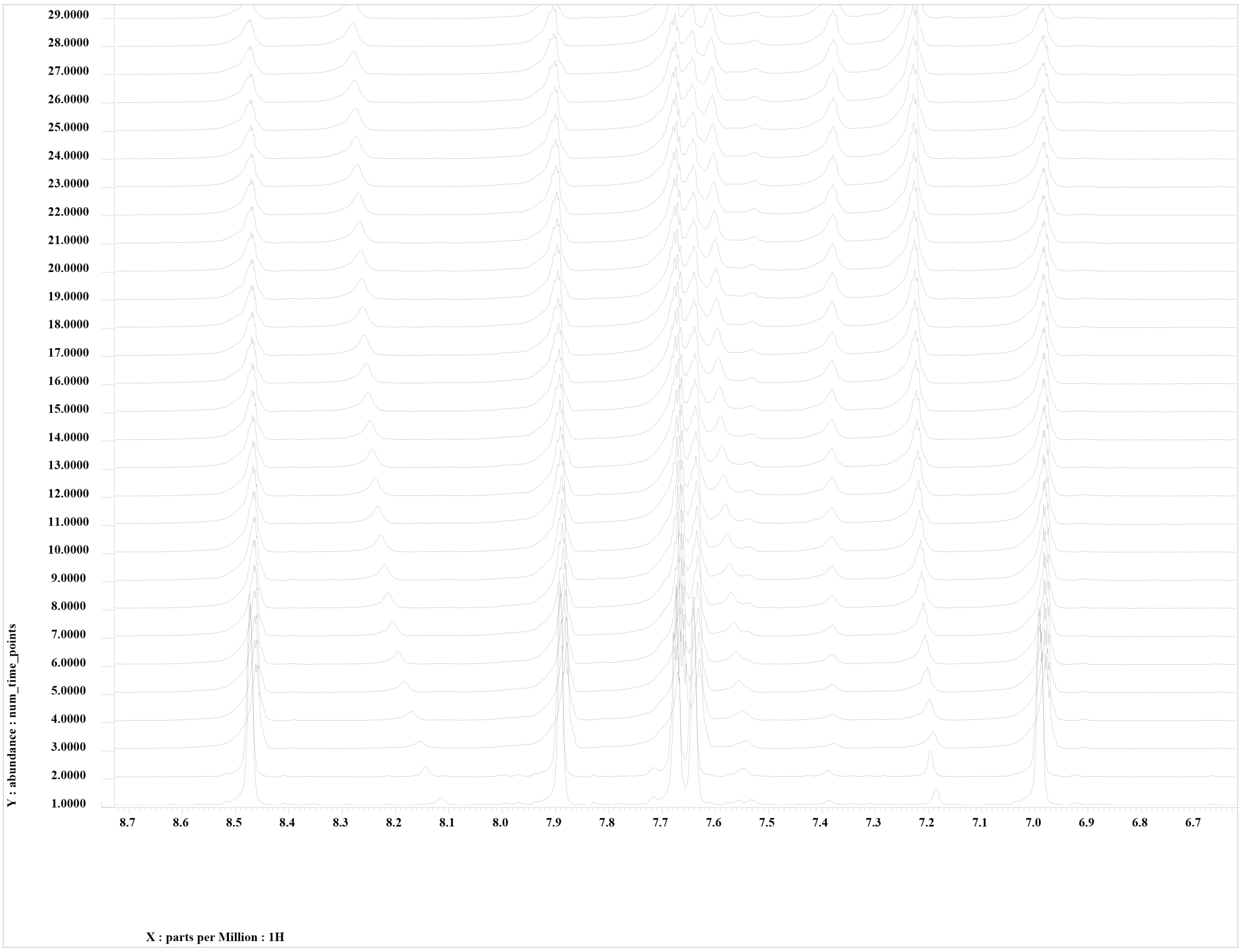


Hydrolysis experiment of 1M4 at 25°C. Showing from 5 minutes to 905 minutes ,where each scan was performed in 30 minutes intervals. Monitoring the disappearance of product peaks at 3.81 ppm and appearance of 3.75 ppm.

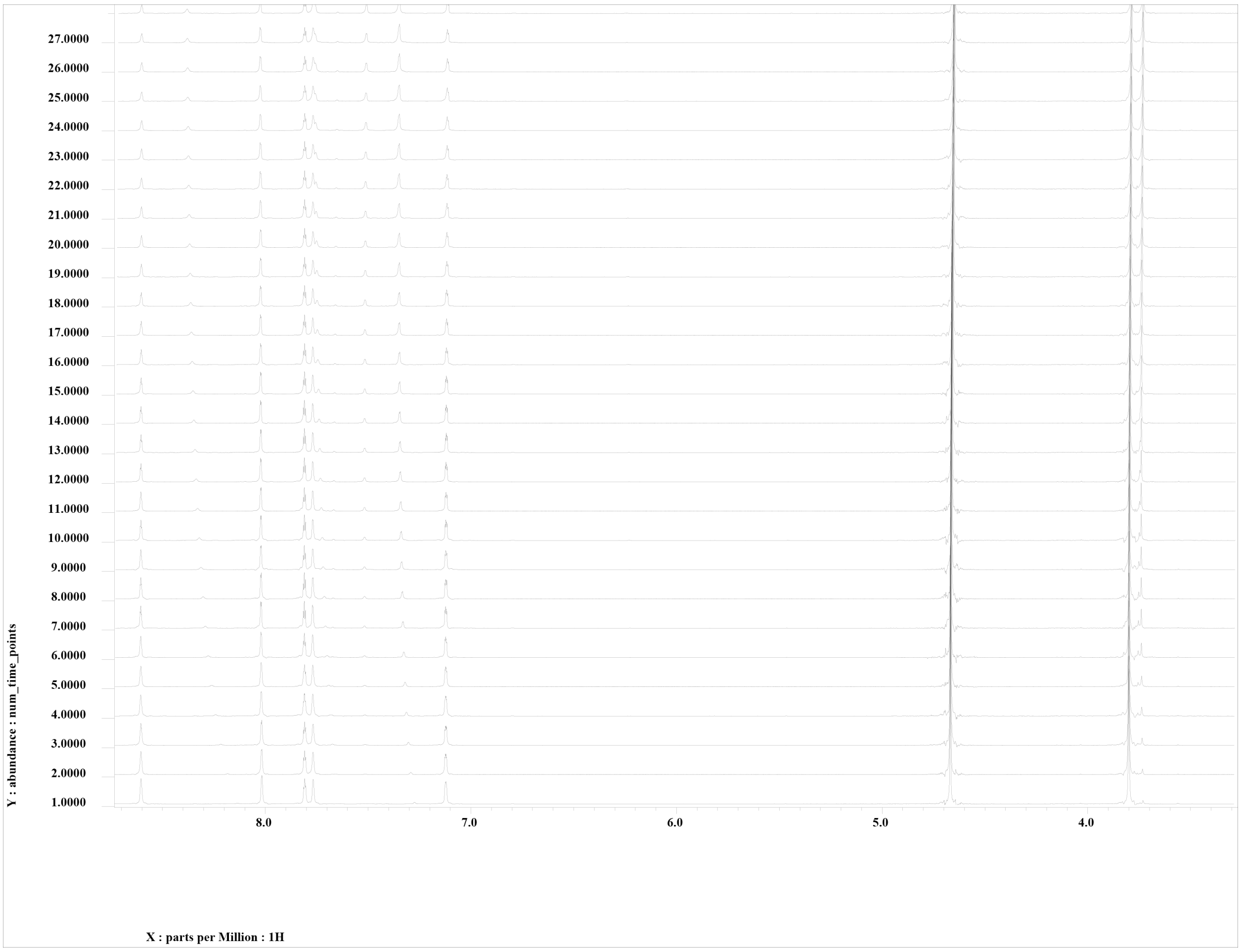


Hydrolysis experiment of 1M4 at 37°C. Showing from 5 minutes to 905 minutes ,where each scan was performed in 30 minutes intervals. Monitoring the disappearance of product peaks at 3.81 ppm and appearance of 3.75 ppm.

**3-azaisatoic anhydride (3AIA)**

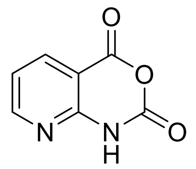

$C_7H_4N_2O_3$, $^{1}H$ NMR (300 MHz, DMSO-d6): δ 7.29 (t, 1H,), 8.31 (dd, 1H), 8.65 (d, 1H), 11.70 (s, 1H) ppm, $^{13}C$ NMR (300 MHz, DMSO-d6): δ 106.71, 119.74, 138.25, 147.12, 153.22, 155.93, 159.63 ppm, HRMS: Compound not identified

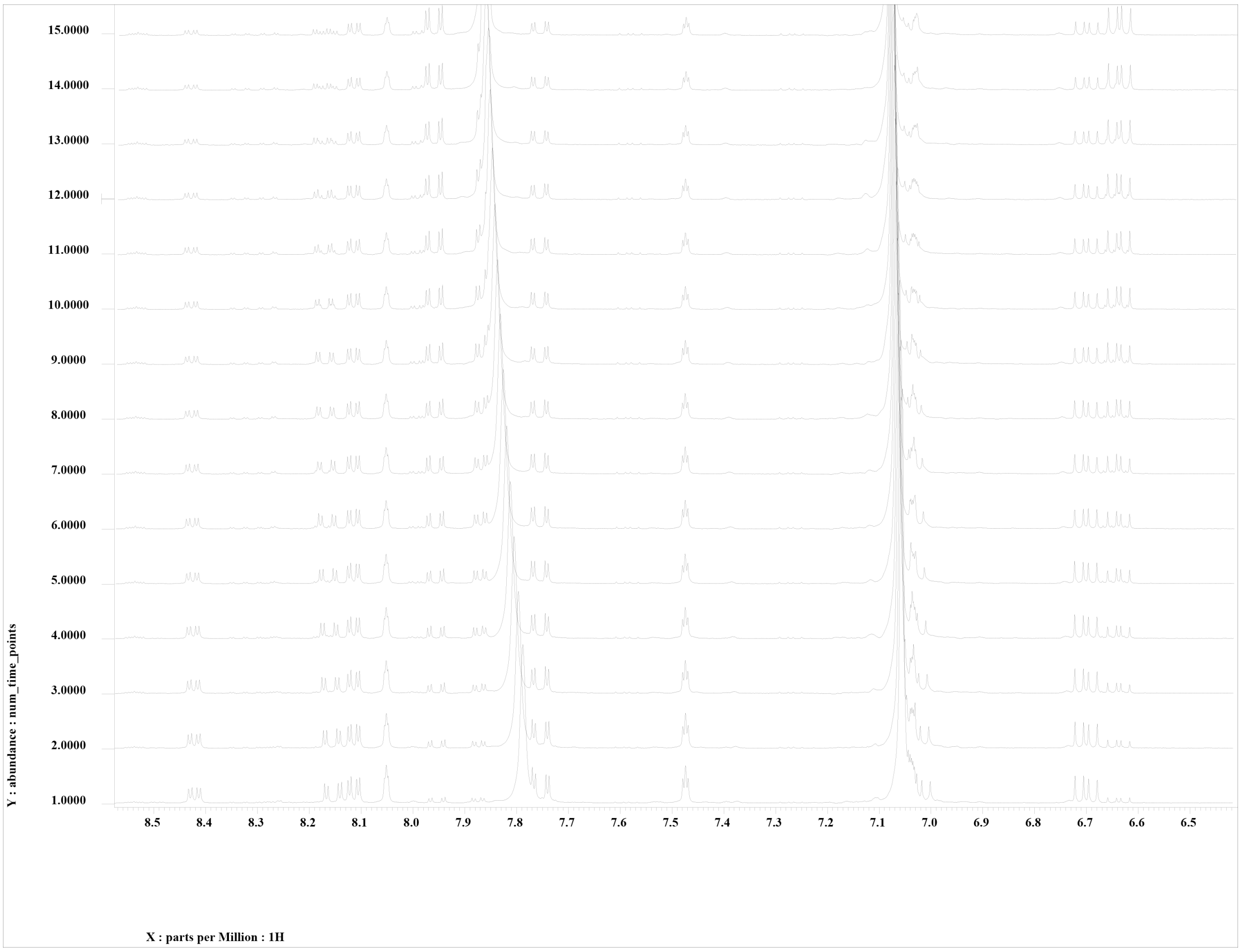


Hydrolysis experiment of 3AIA mixture at 25°C. Showing from 5 minutes to 95 minutes, where each scan was performed in 5 minutes intervals. Monitoring the disappearance of product peaks at 8.42 ppm.

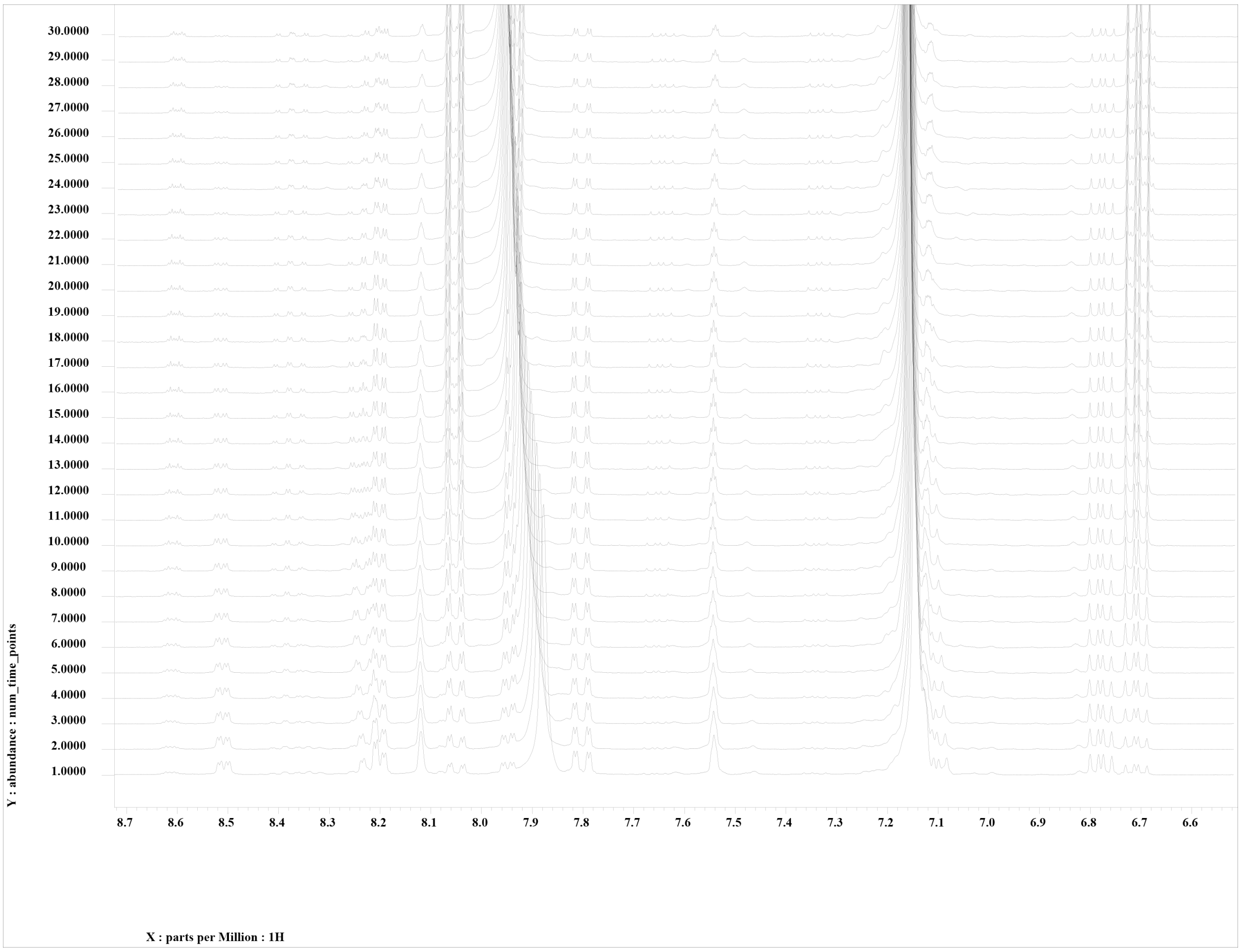


Hydrolysis experiment of 3AIA mixture at 37°C. Showing from 5 minutes to 95 minutes, where each scan was performed in 5 minutes intervals. Monitoring the disappearance of product peaks at 8.42 ppm.
